# Polyadenylated RNA sequencing analysis helps establish a reliable catalog of circular RNAs – a bovine example

**DOI:** 10.1101/2024.04.29.591253

**Authors:** Annie Robic, Frieder Hadlich, Gabriel Costa Monteiro Moreira, Emily Louise Clark, Graham Plastow, Carole Charlier, Christa Kühn

## Abstract

The aim of this study was to compare the circular transcriptome of divergent tissues in order to understand: i) the presence of circular RNAs (circRNAs) that are not exonic circRNAs, i.e. originated from backsplicing involving known exons and, ii) the origin of artificial circRNA (artif_circRNA), i.e. circRNA not generated *in-vivo*. CircRNA identification is mostly an *in-silico* process, and the analysis of data from the BovReg project (https://www.bovreg.eu/) provided an opportunity to explore new ways to identify reliable circRNAs. By considering 117 tissue samples, we characterized 23,926 exonic circRNAs, 337 circRNAs from 273 introns (191 ciRNAs, 146 intron circles), 108 circRNAs from small non-coding genes and nearly 36.6K circRNAs classified as other_circRNAs. We suggested *in-vivo* copying of specific exonic circRNAs by an RNA-dependent RNA polymerase (RdRP) to explain the 20 identified circRNAs with reverse-complement exons. Furthermore, for 63 of those samples we analyzed in parallel data from total-RNAseq (ribosomal RNAs depleted prior to library preparation) with paired mRNAseq (library prepared with poly(A)-selected RNAs). The high number of circRNAs detected in mRNAseq, and the significant number of novel circRNAs, mainly other_circRNAs, led us to consider all circRNAs detected in mRNAseq as artificial. This study provided evidence that there were 189 false entries in the list of exonic circRNAs: 103 artif_circRNAs identified through comparison of total-RNAseq/mRNAseq using two circRNA tools, 26 probable artif_circRNAs, and 65 identified through deep annotation analysis. This study demonstrates the effectiveness of a panel of highly expressed exonic circRNAs (5-8%) in analyzing the diversity of the bovine circular transcriptome.

## Introduction

The current reference genome for cattle (ARS-UCD1.2) is highly contiguous, complete and accurate [1]. The protein coding transcriptome has been well characterized, for multiple different tissues and cell types [2; 3; 4]. In contrast, little is known about how other RNA species are expressed in cattle tissues. In recent years, with the development of improved RNA sequencing methods and bioinformatics tools, to capture and characterize multiple RNA species, circular RNAs (circRNAs), with a closed covalent structure, have emerged as a fascinating new class of RNA molecules. The first two types of circRNAs described in 2012-2013 [5; 6; 7; 8] are now well described and their origin better understood (reviewed in [9; 10]). They are likely to be a natural by-product of the splicing process [11; 12] as other non-co-linear transcripts [13; 14]. During splicing of linear primary transcripts (pre-mRNA), introns (non-coding regions) are spliced out in the form of lariat intronic RNA and exons are spliced together. Classically, a splicing event ligates the 5’ donor site located near the end of the upstream exon (i.e., in the intron on the 3’ side of the exon) with the 3’ acceptor site located near the 5’ side of the downstream exon. The first type of circRNAs is generated by a specific splicing event, known as backsplicing, which results from the splicing of a downstream splice donor to an upstream splice acceptor. For example, at the circular junction we can see the ligation of exon3 end to exon2 start (see M&M_Adoc-1). This backsplicing (BS) leads to an exonic circRNA, which in the vast majority of cases contains only exonic sequences [5; 15; 16]. The genesis of the second type of circRNAs is completely independent of a backsplicing event. Intronic circRNAs contain only intronic sequences and are by-products of classic splicing. The best-known and best-described intronic circRNAs are derived from lariat intronic RNA when intronic lariats escape degradation due to failure of intron debranching. The residue of intronic lariats can become circular RNA precursors to provide ciRNAs or lariat derived circRNAs [8; 17; 18]. In addition to these ciRNAs, intron circles resulting from circularization of the entire intron have also been described [17; 19; 20; 68].

Detection of circRNAs is performed using sequence data from total RNA libraries after depletion of ribosomal RNAs (total-RNAseq) [21]. The identification of a circRNA is always based on the documentation of reads containing the circular junction. Numerous bioinformatic tools are currently available to accurately identify a high number of circRNAs while minimizing the number of false positives [9; 22]. It is important to note that the criteria for managing this balance can vary significantly. A first approach leads to a tool retaining only circRNAs, which meet very precise annotation criteria. The most popular is CIRCexplorer [23], which retains only circRNAs resulting from BS between two known exons (we reserve the term “exonic circRNA” for circRNAs corresponding to this definition) and some putative intronic circRNAs. A second approach to identifying circRNAs is to retain only those suspected of originating from backsplicing; i.e. when the two parts of the canonical splicing motif are found on either side of the interval defined by the circRNA coordinates. This requirement ensures the identification of circRNAs originating from BS and may lead to the identification of circRNAs originating from BS involving unannotated exons. The most popular is CIRI2 [24], which delivers a list of unannotated circRNAs. CIRI2 does not differentiate between exonic circRNAs and putative exonic circRNAs when the putative BS does not involve known exons. An alternative approach is to use a non-restrictive pipeline for the discovery of new types of circRNAs. Liu et al.[25; 26] defined candidate interior circRNAs as those originating from single introns, exons, intergenic regions, and pairs of adjacent introns or adjacent exons (without regular backsplicing). For these interior circRNAs, the presence of repeated sequences appears to be the key to their genesis [26]. To define sub-exonic circRNAs, only circRNAs are retained when both coordinates of the circular junction are located within a single exon [27; 28] even though the class of sub-exonic circRNAs could also be considered as a subclass of interior circRNAs. Several interior circRNAs have been validated using experimental data [25; 26]. The main feature of the interior/sub-exonic circRNAs is that more than one circRNA is detected from the same genomic locus [25; 26; 27; 28].

To manage the balance between true and false circRNAs, it is necessary to have a better understanding of the genesis of false circRNAs. Differentiating between a circRNA observed within a dataset as either generated *in-vivo* or being an artificial circRNA (not generated *in-vivo*) is difficult but necessary [22; 29]. Nielsen et al (2022) points to small circRNAs as particularly suspect [22]. In our previous study [27] we identified several circRNA clusters with unannotated exon boundaries along bovine chromosomes, likely reflecting the presence of sequence/assembly/annotation problems in these regions. The presence of falsely mapped inverted sequences in the genome assembly potentially leads to mapping of reads from the linear transcript as artificial reads spanning the circular junction. Although no circRNA generated *in-vivo* is expected in mRNAseq data (library prepared with poly(A)-selected RNAs), [21; 22], these *in-silico* generated artificial circRNA annotations (*in-silico* artif_circRNAs) would be found in both mRNAseq and total-RNAseq. An analysis including total-RNAseq and mRNAseq prepared from the same sample would be informative to resolve this. A recent study examined two samples to establish the background of the circRNA identification process, but only for exonic and intronic circRNAs [17]. One study has used mRNAseq data to conduct classical circRNAs analyses [30] without indicating that these datasets are a priori unsuitable for characterizing circRNAs [9]. In 2015, Lu et al. reported a comparison performed in rice between the circRNAs detected in mRNAseq and in poly(A)-depleted samples [31]. The total numbers of detected circRNAs in mRNAseq was slightly higher than those in poly(A)-depleted samples. In 2023, Ma et al. [21] suggested that nonspecific binding of circRNAs with oligo(dT)-beads explained the presence of circRNAs in their mRNAseq data. In their study a high fraction of reads were not mapped to the rice reference genome (55% in mRNAseq and 93.4% in poly(A)-depleted), indicating that the data quality may have been low. To resolve this conflict of opinion about the circRNAs present in mRNAseq datasets, we suggested comparing the circRNA content of mRNAseq with that of total-RNAseq from the same samples. This pairwise comparison method has the advantage of providing equal opportunity for any *in-vivo-*generated circRNA to be present in both mRNAseq and total-RNAseq datasets. We noticed there is a similar conflict of opinion regarding datasets generated after RNase-R treatment. This enzyme is used to eliminate the majority of linear transcripts and increase the concentration of circRNAs [22]. Some authors consider circRNAs detected only after treatment to be low-expressed circRNAs [5], while others do not consider them as reliable circRNAs [32].

One of the aims of the European BovReg project (https://www.bovreg.eu/) was to generate a map of functionally active regulatory and structural elements in the bovine genome using a diverse catalog of at least 26 tissue types collected from individuals of both sexes and from divergent breeds/crosses (117 samples in total) [4; 33]. The data generated by BovReg provided an interesting opportunity to explore some aspects of the circRNA transcriptome in cattle because the transcriptome sequencing was performed in two ways: mRNAseq and total-RNAseq. The respective datasets were generated in very similar conditions to minimize any batch effects, and paired mRNAseq and total-RNAseq datasets were available for a subset tissues from the same animals. We performed the characterization of circRNAs using two bovine annotations (the current Ensembl version 105 and a new annotation generated by the BovReg project) and 117 samples obtained from 26 tissues, across 3 populations of cattle, with a first objective to understand the presence of non-exonic circRNAs. For this purpose, we also looked at a subset of 63 samples with available high-quality paired mRNAseq and total-RNAseq data. In a previous study performed on bovine, ovine and porcine tissues [27], we had already obtained an indication of a large proportion (40 to 80%) of non-exonic/intronic circRNAs in bovine and ovine tissues. In this current study, we again could only annotate 40% of the highlighted circRNAs as exonic circRNAs in spite of a more comprehensive transcriptome annotation. With the availability of paired datasets, we had the chance to further explore mRNAseq-based output with respect to circRNAs. By performing these analyses, however, we did not expect to fully resolve the question that arose regarding the low proportion of circRNAs annotated as exonic circRNA. In this study, firstly we aimed: i) to understand the presence of non-exonic circRNAs in the cattle transcriptome, ii) to understand the origin of artificial circRNA, i.e. circRNA not generated *in-vivo*. Our second objective was to perform a comparison between the circular transcriptome (by considering only reliable circRNAs) of divergent tissues. Our results reflect the diversity of the circular transcriptome in cattle and provide a resource for comparative analysis across cattle populations and between species.

## Materials and Methods

### Animals, samples and datasets

The six animals chosen for the sample collection originated from three populations kept in different environments representing different ages and sexes. Holstein Friesian calves from Belgium (neonatal: male calf 24 days and female calf 22 days), Kinsella composite juveniles from Canada (bullock 217 days and heifer 210 days) and Charolais x Holstein F2 cow and bull from Germany (adult: bull 18 months and cow 3 years, 7 months and 13 days).

All details of the animals are available in [4]. Details of tissue sampling and storage, and RNA extraction, quality and integrity assessment are described in [4]. All samples were sequenced in two ways: mRNAseq and total-RNAseq libraries were generated, quantified and sequenced by the GIGA Genomics platform (University of Liège, Belgium). mRNAseq libraries were built using the ‘TruSeq Stranded mRNA Library Prep’ kit (Illumina) following the protocol provided by the manufacturer. Total RNA libraries were built using the ‘TruSeq Stranded Total RNA Library Prep Gold’ kit (Illumina) following the protocol provided by the manufacturer. The Illumina NovaSeq 6000 instrument was used for sequencing, with a paired-end (PE) protocol (2×150bp).

For circRNA characterization, we considered a batch of 117 datasets obtained by total-RNAseq. Among the 26 tissues represented in this batch, 11 and 4 tissues were represented by 6 and 5 datasets, respectively, see STab-1. A sub-batch of 63 samples were chosen for the comparison of total-RNAseq (63T) and mRNAseq (63m). Among the 11 tissues represented in this batch, 8 and 3 tissues were represented by 6 and 5 datasets, respectively, see STab-1.

### Circular RNA detection and characterization

The RNAseq reads were mapped to the bovine genome reference assembly ARS-UCD1.2 (GCA_002263795.2) using the rapid splice-aware read mapper Spliced Transcripts Alignment to a Reference (STAR, version 2.7.10a [34]). We selected the single-end alignments mode of STAR (STAR-SE) mapping mates of each pair independently. STAR was used with the previously proposed parameters [35] that enable highlighting chimeric reads with only two mapped segments and with a minimal size for the smallest mapped segment of 15 bp.

Our approach to characterize circRNAs is described in Figure 1 (see M&M_Adoc-1-4 for details). The chosen circRNA tool, CD (CircDetector, [27; 36]), works in two steps. The initial step involves identifying reads that contain a circular junction, referred to as circular chimeric reads (CCRs), and generating two output files (Figure 1). In the main CD_detection output file, detection.bed, CD reports a list of all circular RNAs and their associated number of CCRs, each circular RNA being defined by the coordinates of the circular junction (chromosome:start-end|strand). When CD is used with a gtf_file containing exon features, the second module of CD is able to annotate certain circRNAs (Figure 1, see also M&M_Adoc-2 for details). For instance, CD can identify circRNAs resulting from backsplicing events and provide a list of putative sense-exonic circRNAs. It also identifies the two exons involved in the backsplicing and their respective parental gene. (Figure 1). In this study, we defined other_circRNAs as those not included in any of the three retained lists (Figure 1). By creating this category, we set these circRNAs aside with the *de-facto* suspicion that some of them are artif_circRNAs, to be examined in detail in this study. We performed manual curation of each of the three retained output files to identify exonic circRNAs (associated with the use of blue color in the figures), lariat-derived circRNAs (black/yellow), intron circles (black/pink), and sub-exonic circRNAs from small non-coding genes (snc) (black/green). For example, we rejected a potential exonic circRNA candidate, which would have resulted from backsplicing between two exons from different parental genes (Ensembl_gene A and Ensembl_gene B; or MSTRG_gene X and MSTRG_geneY). CircRNAs that were rejected during manual curation (symbolized by a trashcan in Figure 1) were not added to the other_circRNAs list (orange in the figures).

**Figure 1:**
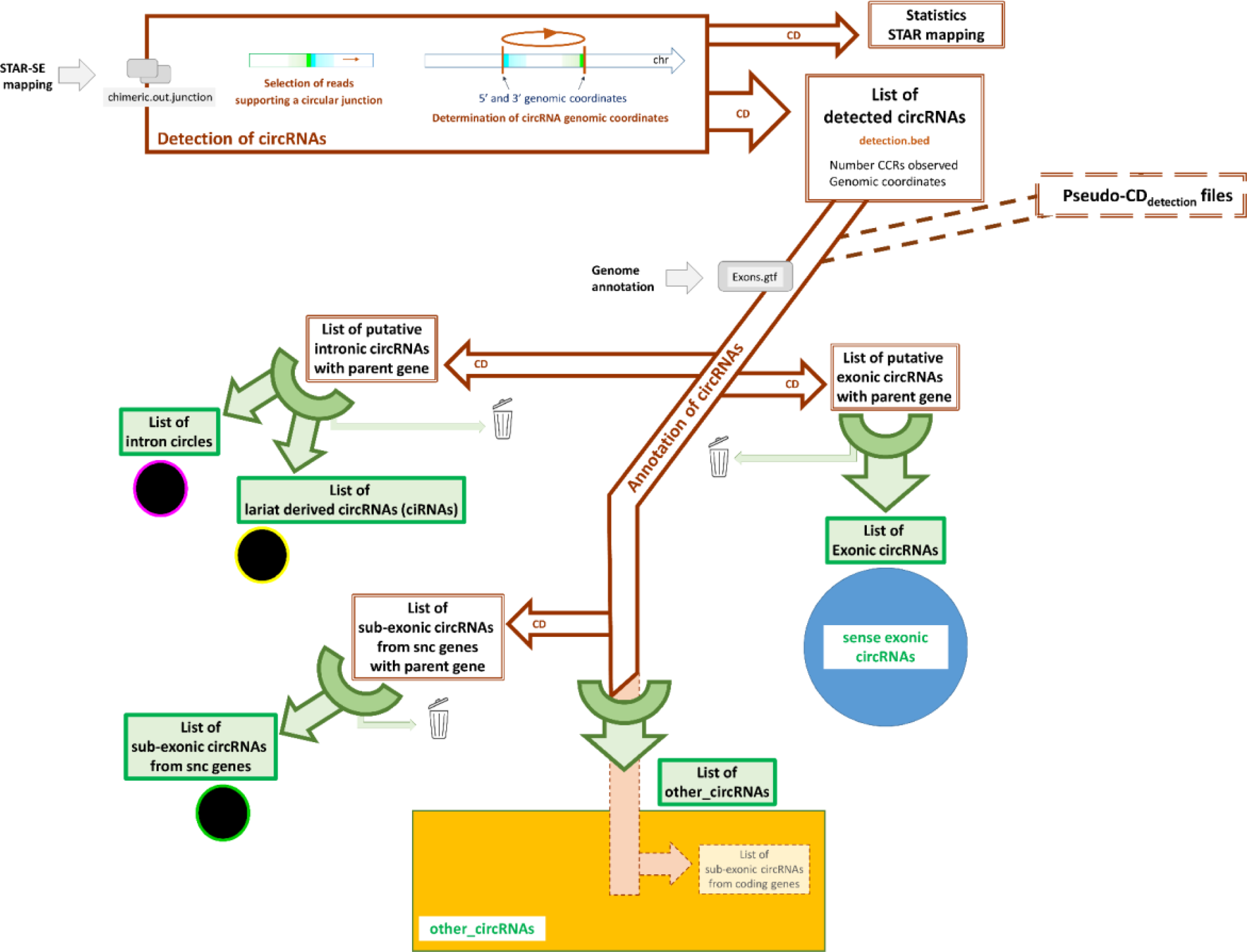
Analytical pipeline used to characterize circRNAs. This pipeline combines the uses of the CircDetector (represented in brown) in two steps and several small manual process (represented in green). The input files for the CD are represented in gray frames, while the output files are represented in double brown frames. The first part of this pipeline is managed exclusively by CD, and is shown horizontally at the top. For this detection step, users can select parameters to exclude sporadic circularization events and loci that are too small to be reliable circRNAs. In addition, of the main CDdetection output file (detection.bed), CD produces a second file reporting all statistics of STAR mapping. In a second step, CD is able to identify several types of circRNAs. In our approach, we retained three lists provided by CD (exonic circRNAs, intronic circRNAs and a list of sub-exonic circRNAs deriving from genes identified in the gtf_file as small non-coding (snc) RNA). We defined other_circRNAs as those not included in any of these three lists. The lists of circRNAs retained after manual verification are shown in green rectangles. The circRNAs excluded by this manual curation do not join the other_circRNAs list, but are declassified (symbolized by a trashcan). For more details and examples, see M&M_Adoc-1-4. The source code of the CD is available from https://github.com/GenEpi-GenPhySE/circRNA.git.

Annotating a large number of circRNA lists has never been performed previously as an individual task. This step was performed with pseudo-CDdetection output files using in particular a compilation of all detection.bed files from all datasets (Figure 1). To identify antisense exonic circular RNAs (AS-exonic circRNAs), we generated a new pseudo-CDdetection output file by reversing the strand of each circRNA. Subsequently, we executed the annotation step again, resulting in a comprehensive list of exonic circRNAs, which enables access to the corresponding list of AS-exonic circRNAs.

In addition, detection of circRNAs was also performed with CIRI2 [24]. Unlike CD and most other circRNA discovery tools, CIRI2 works from paired ends alignments. It requires these alignments to be performed by BWA-MEM (here we used the version 0.7.17-r1188) [37]. CIRI2 implements multiple filtering strategies to eliminate false positives, including splice site analysis. CIRI2 (version 2.0.6.) was used with default parameters and only circRNAs detected by two reads (the maximum threshold option provided by the program) containing a circular junction were retained. We have chosen to use CIRI2 without an annotation file and therefore the output file does not include any information about the parental gene. This output file contains the number of reads spanning the circular junction. The annotation of CIRI-circRNAs was performed by using CD with a pseudo-CDdetection output file. We used the BovReg annotation for classifying exonic and intronic circRNAs. For exonic circRNAs, we did not add a manual verification process; as a result, we identified only putative-exonic circRNAs.

### Analyses relative to exonic sequences

The BovReg annotation consisted of a gtf_file defining 683,396 distinct exons (average length = 1,628 nt and median length = 226 nt). Only 235,049 were previously described by Ensembl v105 (average length = 308 nt and median length = 139 nt).

To perform what we call a minimal_annotation of exonic circRNAs, we built two sub-lists (Left_exons and Right_exons) from the list of all BovReg exons. To constitute the list of Left_exons, we selected exons according to their unique first genomic coordinate (M&M_Adoc-4) keeping only the exon with the smallest size in case of multiple exons with the same first coordinate (M&M_Adoc-5A). For the list of Right_exons, we only filtered for unique second coordinates (M&M_Adoc-5A). We retained a list of 636,307 distinct exons (582,688 in the list of Left_exons and 456,432 in the list of Right_exons). To perform a manual and “minimal” annotation of exonic circRNAs, it is necessary to identify the two exons involved in backsplicing using these two lists (M&M_Adoc-5B). The list of Left_exons is used to identify the upstream exon (or “acceptor exon”) involved in the backsplicing when the parental gene is located on the forward strand and the downstream exon (or “donor exon”) when the parental gene is on reverse strand (see M&M_Adoc-1).

We used several tools available on the Galaxy platform proposed by Sigenae [38] in particular to perform *bedintersect* (http://bedtools.readthedocs.io/en/latest/content/bedtools-suite.html). To examine the exonic sequence content of the genomic interval defined by the circRNA, we applied a 90% exon overlap threshold on the same strand. This allowed us to conclude that the circRNA interval contained what we then call a *quasi-full exon*. *Bedintersect* was also used to analyze the localization of the points defined by the two genomic coordinates of a circRNA, both inside and outside of an exon. The search was performed using the BovReg list of 683,396 exons, considering both strands. For each circRNA, two genomic intervals were defined: the first interval contains the 30 nucleotides downstream of the 5’ coordinate, and the second interval contains the 30 nucleotides upstream of the 3’ coordinate.

### Statistical analyses

All the statistical analyses were carried out using R (v.4.0.2) [39]. Significant differences between circRNA proportions from contingency tables were identified with the Pearson’s Chi-squared test (chisq.test function from R STAT package v.4.0.2) [39]. A p-value less than 0.05 was considered as statistically significant.

### Hierarchical clustering and principal component analyses

The hierarchical clustering analyses (HCA) were performed on the Galaxy platform proposed by Sigenae [38]. All clusters were done with the “ward” agglomeration method as suggested by developers [40] and using Pearson’s correlations as distance. The principal component analyses (PCA) were also performed on this platform, with the function PCAFactoMineR, using the FactoMineR package. Data was transformed by the normalization module available on the Galaxy Platform. For HCA, the log-binary (binary log (expr + 0.0001)) and standard score (standard score; mean=0 - sd=1) methods were used, while for PCA only the standard score method was used. These tools are part of a set of statistical tools made available by members of the BIOS4BIOL group (”Normalization”, “Summary statistics”, “Hierarchical clustering” and “PCAFactoMineR”) (see https://github.com/Bios4Biol).

For clustering, 96 samples were retained (see STab-1). To avoid introducing a tissue represented by a single dataset, we selected 15 tissues, where samples were available for the two youngest (neonatal) and at least three of the older animals (juvenile or adult). In addition, we considered five tissues, where samples were available for the two young animals. For the PCA analysis of samples related to reproduction and hormonal function, 19 samples were considered from pituitary gland, adrenal gland, ovary, testis, uterus, and uterus-horn. For more details, see STab-1.

## Results

### Characterization of 61,083 circRNAs less than 40% of them being exonic circRNAs

For circRNA characterization, we retained 117 samples sequenced with total-RNAseq from 26 distinct tissues providing a total of 10,052 million reads (150 bp) across all the samples. Three tissues samples from neonate animals (jejunum-female, rumen-male, pancreas-male) were sequenced at high depth (called “XL sequencing”). Two sequencing datasets were available for the cerebral cortex sample from the juvenile castrated male. 86% of reads mapped unambiguously to the bovine reference genome. At least 37 million uniquely mapped reads were obtained for each sample. For the three datasets with XL sequencing, 395, 410 and 432 million reads were available, while for the 114 other datasets we observed an average of 77 million reads that were uniquely mapped (all details are available in STab-1). We did not observe any outlier samples, with a poor mappability, and all 117 datasets were considered for further analyses.

We started with the exhaustive list of circRNAs present in at least one sample. Rare circularization events were excluded by only retaining circRNAs detected by five reads containing the circular junction (CCRs for circular chimeric reads). Several studies have demonstrated the value of excluding such events [32; 41]. The 117 output files generated by CD [27; 36] were concatenated, resulting in the detection of 66,299 circRNAs. After discarding circRNAs with genomic size less than 70 nucleotides 61,083 circRNAs were retained.

Annotation with CD was performed using the list of 61,083 circRNAs as a single pseudo-sample using the two bovine annotations. All selected output files were manually inspected (Figure 1, see also M&M_Adoc-6). We were aware that the list of circRNAs provided by CD can include false entries and we chose to retain only three sub-lists created by CD (see materials and methods, and Figure 1) and put the others in a list of other_circRNA. To create this list, we deducted from the initial list of 61,083 circRNAs the following: 24,359 putative sense-exonic circRNAs (CD BovReg annotation), 373 putative intronic circRNAs (CD Ensembl annotation), and 108 sub-exonic circRNAs from snc genes (CD Ensembl annotation). Therefore, we retained 36,215 circRNAs as other_circRNAs for further analysis (Figure 2A1).

**Figure 2:**
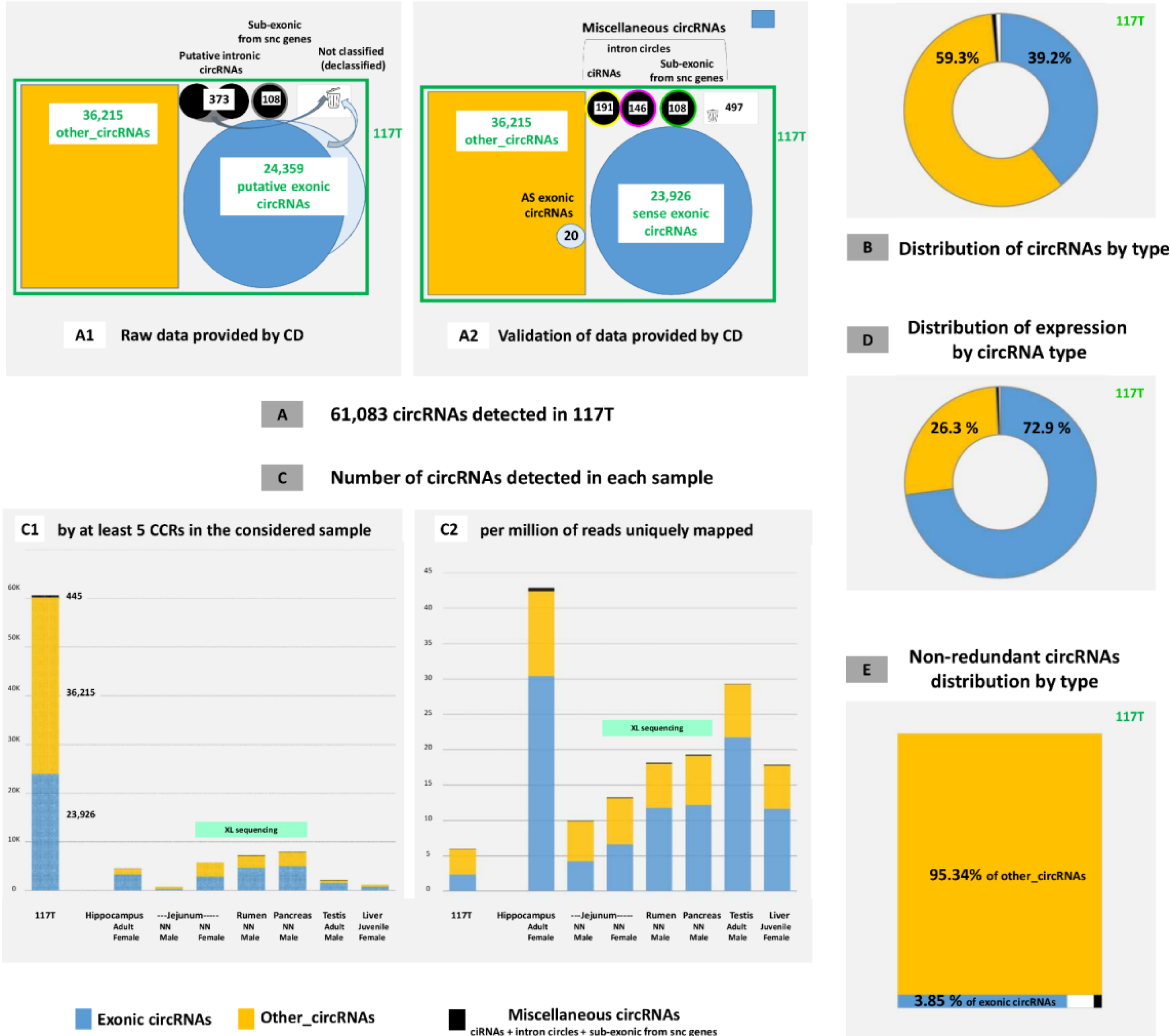
Overview of circRNAs detected in the 117 samples considered. **(A**) circRNAs detected in 117T (symbolized by the green frame) **(A1)** 61,083 circRNAs were retained after the characterization by CD (with a minimum genomic size of 70 nt and whose presence has been attested by at least 5 reads supporting the circular junction in at least one sample). After the examination of the annotation suggested by CD, we put in a new category, other_circRNAs, 36,215 circRNAs for further analysis symbolized by the orange rectangle) (see also Res_Adoc-10). **(A2)** We retained 23,926 sense-exonic circRNAs (blue disk), 191 ciRNAs, 146 intron circles and 108 sub-exonic circRNAs from snc genes (represented by three black discs). We identified also 20 AS-exonic circRNAs, represented by a light blue disc inside the orange rectangle of other_circRNAs since the 20 identified fell into this category. **(B)** Distribution of the 61,083 circRNAs by type. **(C)** Number of circular RNAs. The first histograms at the left concern the full-virtual sample, named 117T. The other seven histograms consider data from six individual samples from six different tissues. NN=neonate. The three deeply sequenced samples are marked with a green label “XL sequencing” above the histograms. **(C1)** Number of circRNAs validated by the detection of 5 CCRs in the considered sample. **(C2)** Number of circRNAs validated per million reads uniquely mapped. **(D)** Distribution of expression based on circRNA type. **(E)** Distribution of the 6,982 non-redundant circRNAs by type.

The analysis of the list of putative sense-exonic circRNAs led to the characterization of 23,926 sense-exonic circRNAs (i.e. resulting from backsplicing (BS) involving known exons; Figure 2A2). More precisely, we were only able to annotate 20K circRNAs as exonic circRNAs using the Ensembl annotation and to add 4K circRNAs to the list of exonic circRNAs using the BovReg annotation. They are originating from approximately 8K parental genes (Res_Adoc-1). Furthermore, we observed 2K exonic circRNAs from a backsplicing between an exon with an Ensembl ID and a novel (MSTRG) exon in the BovReg annotation. For example, in the region of *SMARCA5* (BTA17: 14,342-14,380 Mb), eight circRNAs were identified and seven as sense-exonic circRNAs (Figure 3). From the sub-list of putative intronic circRNAs provided by CD, our analyses featured the annotation of 191 ciRNAs (147 introns concerned from 146 genes) and 146 intron circles (126 introns concerned from 124 genes) (list in STab-2). We have grouped circRNAs that do not fit into the two main categories (sense-exonic circRNAs and other_circRNAs) into “miscellaneous circRNAs” (Figure 2A). We found ciRNAs, intron circles and sub-exonic circRNAs from snc genes. In these 117 samples considered, 39.2% of circRNAs are exonic circRNA and we classified 59.3% as other_circRNAs (Figure 2B).

**Figure 3:**
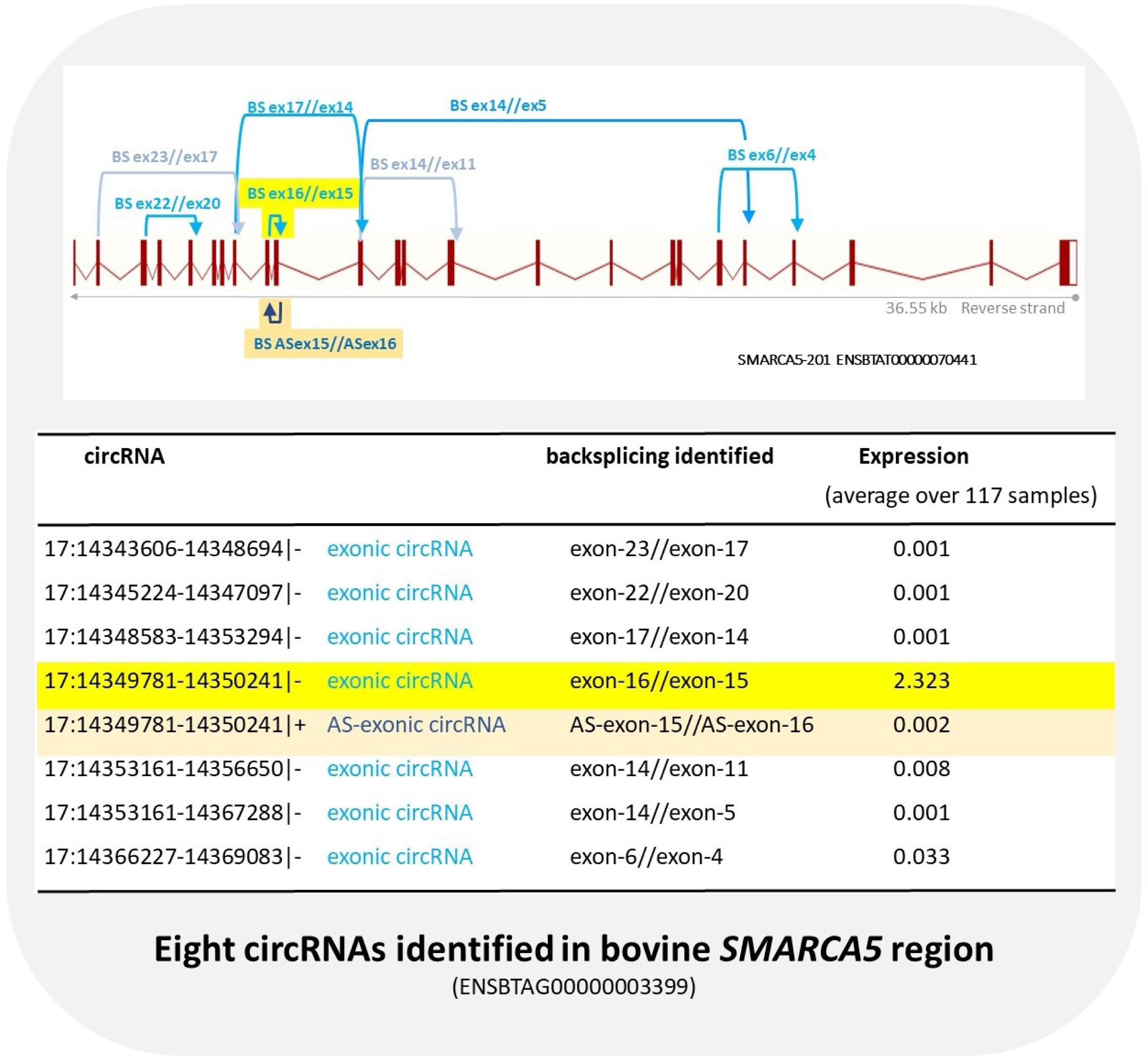
circRNAs from the *SMARCA5* region. This figure shows the bovine region of *SMARCA5* (ENSBTAG00000003399; BTA17: 14,342-14,380 Mb). Eight circRNAs were characterized with seven sense-exonic (BS in turquoise) and one antisense-exonic (BS in navy blue) circRNAs. As is often the case, the antisense (underlined in orange) has identical coordinates (except for the strand) to the most highly expressed sense-exonic circRNA (underlined in yellow). This exonic circRNA resulting of a BS between the exons 16 and 15 (BS ex16//ex15; 17:14349781-14350241|+) was also identified in human and sheep. Figure constructed from data and drawing uploaded on July 27, 2023 from the Ensembl bovine genome database (http://www.ensembl.org/Bos_taurus/Info/Index).

### Analysis of circRNA diversity observed in 117 bovine datasets (total-RNAseq)

In the full pseudo-sample (117T), we observed 61,083 circRNAs (Figures 2A2 and 2C1), with an average scaled read count of 6 circRNAs per million of uniquely mapped reads (Figure 2C2). The landscape of the 117T merged list is very different from the 117 individual samples (Figure 2C). We observed less circRNAs per million of uniquely mapped reads in 117T than in each individual sample (Figure 2C2): however, while the number mapped reads add up across the samples, the number of distinct circRNAs does not due to redundancy of circRNA identification in the different samples. The seven examples in Figure 2C show the great diversity of these 117 individual datasets. Three samples had very deep sequencing (XL sequencing), and this led to the identification of more circRNAs, including predominantly more of the other_circRNAs type (Figure 2C1).

We observed that 6,982 circRNAs were detected in a single sample and were not confirmed in any other samples, not even when applying a threshold of a single CCR. It is hardly surprising to find the three samples that benefited from XL sequencing were among the samples with the highest proportion of circRNA not found in any other sample. Among the 6,982 non-redundant circRNAs detected in 117 datasets, only 268 were exonic circRNAs, i.e. we have 95.3% of other_circRNAs (Figure 2E). In other words, more than 18% (6,714 out of 36,215) of the other_circRNAs were detected by only five CCRs and in only one sample.

### Brief description of CD-other_circRNAs

When we looked at the size of the genomic interval defined by the two genomic coordinates of an other_circRNA, we noted that 22.9% of other_circRNAs defined small genomic intervals (see Res_Adoc-2). When we looked at the exon content of the genomic interval defined by the two boundaries of the other_circRNAs, we found that 71.2% did not contain a quasi-full exon (90% of an exon, see MM section) (see Res_Adoc-3). Nevertheless, 62.4% of the other_circRNAs have their two genomic boundaries in exonic sequences (see Res_Adoc-3, and M&M_Adoc-3).

Among the 61,083 circRNAs, we identified 487 from the mitochondrial genome, all included in other-circRNAs. Other_circRNAs from the mitochondrial genome are more often smaller than other_circRNAs detected on the nuclear chromosomes (data reported in Res_Adoc-2 showed a statistically significant difference). As in our previous study [27], we identified several clusters of other_circRNAs along the chromosomes, likely reflecting the presence of sequence/assembly/annotation problems in these regions. We identified 3,187 circRNAs (including 3,159 other_circRNAs) clustered in a region (BTA-27: 6.21-6.23Mb) known to contain the *Defensin* gene. Other_circRNAs from the *Defensin* region are less often of small size than other_circRNAs detected on other chromosomes (data reported in Res_Adoc-2 showed a statistically significant difference). The assembly of this region is thought to be difficult due to a substantial number of copies of the same or very similar sequences. In addition, it is assumed that bovine individuals differ in the number of *Defensin* gene copies. As such, this region is clearly a candidate to provide *in-silico* artif_circRNAs.

### Brief description of circRNA expression

The number of circular junction reads associated with the detection of a circRNA is commonly used to quantify the expression of that circRNA (corrected for the number of uniquely mapped reads in the dataset). To determine the expression of each circRNA in each sample, an inventory integrating statistic relative to each sample was obtained from a second run (117X) of CD, but without a threshold on the number of CCRs (see also M&M_Adoc-6). At expression level, we noted a clear dominant impact of exonic circRNAs, since they are responsible for 72.9% of the global expression of circRNAs in our datasets (Figure 2D). This result is in contrast to the 39.2% of circRNAs annotated as exonic circRNAs (Figure 2B). Intronic circRNAs are responsible for 0.37% of the global expression of circRNAs (Figure 2D). The expression of intronic circRNAs, ciRNA-ATXN2L, which was found to be the dominating ciRNA in pigs [28], was very low in most samples in the current analysis of bovine tissues, similar to a previous report [36].

Although the expression of an exonic circRNA varied between 0 and 30, we defined “notable expression” as expression above 0.05. We observed that 95.5% (22,846/23,926) of exonic circRNAs had a notable expression in at least one sample but only 5.3% (1,268/23,926) had a notable expression on average across all 117 samples. For example, eight circRNAs were identified in the region of *SMARCA5* (ENSBTAG0000000003399), but only one has a notable expression on average across all 117 samples (Figure 3). This exonic circRNA (17:14349781-14350241|-) has a notable expression in each of these 117 samples (STab-3).

### Highlighting of particular exonic circRNAs

#### Identification of 20 antisense exonic circRNAs

In addition, among the 23,926 sense-exonic circRNAs, we characterized 20 that seemed to be the result of a backsplicing between two exons of the parental gene considered in antisense (AS-exons). Among all the reverse-complement sequences of known exons (AS-exons), we searched for those involved in antisense-backsplicing (AS-BS). We chose a methodology that proved effective even when the exons were described on only one strand. In the STab-4, we reported the expression of all sense and antisense exonic circRNAs for 20 regions. When an AS-exonic circRNA was observed (M&M_Adoc-3, circRNA-2), the corresponding sense-exonic circRNA was detected in 19 cases out of the 20 considered. In 13/19 cases, the corresponding sense-exonic circRNA is the exonic circRNA with the highest expression among all circRNAs produced by this gene. In addition, the AS-exonic circRNA always had a very low expression. All antisense-exonic circRNAs were found in the list of other_circRNAs (Figure 2A2). Figure 3 shows an example (*SMARCA5*) selected from these 20 regions. The characterization of these 20 AS-exonic circRNAs, suggested the existence of 40 AS-exons really used in AS-BS. Among the 683,396 described exons in the bovine genome by the BovReg annotation, we detected 145, which were described in the forward strand and in the reverse strand with the same genomic coordinates. These two lists of 40 and 145 exons have no exons in common. The discovery of new exons involved in these 20 AS-exonic circRNAs explains why they have been found among the other_circRNAs.

#### Focus on three exonic circRNAs (from TTN, SMARCA5, and BTBD7)

It is interesting to highlight three particular exonic circRNAs (see also STab-3). (1) An exonic circRNA with expression restricted to a single tissue (heart). This circRNA (2:18153915-18180018|+) originates from a region very poorly annotated in Ensembl, probably harboring the *TTN* region. Nevertheless, the list of exonic circRNAs (see STab-3) from this region is long but it is unique with this tissue specificity (only detected in neonates Belgian animals). (2) The circRNA with the second highest average expression across the 117 samples was a circRNA from *SMARCA5* (BS ex16//ex15; 17:14349781-14350241|-; ENSBTAG00000003399). In addition, it was detected and validated in all 117 samples. We also detected the corresponding antisense exonic circRNA. This gene (35 exons, 36.55 kb) is capable of producing six other sense-exonic circRNAs (Figure 3). The exonic circRNA resulting of a BS between the exons 16 and 15 (BS ex16//ex15; 17:14349781-14350241|+) was also identified in human (its name is *circSMARCA5(15,16)* according to the nomenclature proposed by Chen et al. 2021 [42]) and studied very extensively [43]. It was also detected in sheep tissues [27]. In contrast, it has never been detected in pigs [27]. Instead, in pigs we previously detected ExoCirc-9244 (created by exon 16-exon 12 backsplicing of the *SMARCA5* linear transcript) at high frequency, but only in the testis at a specific stage of puberty [17]. No exonic circRNA resulting from BS ex16//ex12 are produced in cattle. This is a good illustration that the ability to produce exonic circRNAs can be conserved in four species, but exonic circRNA is not systematically orthologous [32]. (3) The recently published study of an exonic circRNA from the bovine *BTBD7* gene led us to focus on this gene with 11 exons and a length of 85.6 kb (ENSBTAG00000046185) [44]. We have characterized only one exonic circRNA produced by this gene. This is the one characterized by Ma et al. (2023) and it was detected in 117/117 samples. Although we observed that the highest expression of this exonic circRNA is not found in subcutaneous fat, our observations on tissue expression of this exonic circRNA are almost all compatible with the results published by Ma et al. (Res_Adoc-4).

#### Emphasis on the remarkable production of exonic circRNAs by DOCK1and NEB

Eight exonic circRNA are produced from *SMARCA5,* which is a gene with 25 exons and a length of 36.55 kb (Figure 3). Therefore, it is interesting to focus on two very large genes able to produce several exonic circRNAs. Our first choice is *DOCK1* (ENSBTAG00000031890, 52 exons, 556 kb) because the mechanism of biogenesis of a particular exonic circRNA produced by human *DOCK1* (circ*DOCK1*(2,27)) has recently been studied [45]. In the bovine region, 24 circRNAs were detected, all annotated as exonic circRNAs, and 116/117 samples were affected by non-null expression of an exonic circRNA from *DOCK1*. Even though the exon-2 is involved in the BS for five exonic circRNAs, the BS between the exon 27 and the exon 2 was not observed in bovine. Our second example is *NEB* ((ENSBTAG00000006907, *Nebulin*, >160 exons, 219 kb) because a recent study was published on a specific exonic circRNA from this bovine gene [46]. The very peculiar structure of the major transcript led Huang et al. to suggest that this exonic circRNA may encode a rolling-translated peptide that is highly homologous to a portion of the NEB protein. In this study, 14 exonic circRNAs from *NEB* were characterized and detected almost exclusively in muscle samples. However, the exonic circRNA from *NEB* characterized by Huang et al. was not found and no possible surrogate was identified. Due to the limited size of the muscle samples in this study (6/117), we also examined the list of circRNAs characterized by CircExplorer2+CIRI2 in a study of 12 muscle samples from adult animals (and 15 liver and 6 testis samples) [27]. We found 25 novel *NEB* exonic circRNAs, including three exonic circRNAs compatible with a rolling translation peptide, but not the one identified by Huang et al (see for details Res_Adoc-5). In contrast to *SMARCA5* (Figure 3, 8 exonic circRNAs), we are not able to distinguish among the 24 exonic circRNAs from *DOCK1* or among the 14 from *NEB* one exonic circRNA that dominates the others by its expression (STab-3). The examples of *DOCK1* and *NEB* illustrate the well-known fact that genes with a large number of exons are often able to produce a wide range of exonic circRNAs [17; 47]. We suspect that there may be poor reproducibility (across species, animals, or tissues) in this circRNA production. This study also highlights that for these genes, the priority might be circRNA production to limit parental gene expression, rather than the exact exon content of the produced circRNAs.[32; 48].

#### Imperfect backsplicing for exonic circRNAs

An example to demonstrate the existence of imperfect backsplicing and its consequences on the circRNA list is provided for the *MORC3* (ENSBTAG0000000011876) gene. Among the circRNAs detected in this region by CD, we focused (Figure 4A) on seven other_circRNAs and one exonic circRNA (BS exon7//exon5, 1:148622260-148626199|+, Figure 4A) that share their downstream boundary (on the donor side, here the end of exon7). No tissue specificity was observed for these eight circRNAs. They may all be present together. For example, all eight were observed in the sample of subcutaneous fat, thyroid, uterus and adrenal glands from the 9-month-old female. The two most common (Figure 4A, 1:148622261-148626199|+ and 1:148622265-148626199|+) were detected in all 117 samples by CD. Three of the seven other_circRNAs from *MORC3* are also detected by CIRI2 (Figure 4A, 1:148622241-148626199|+; 1:148622265-148626199|+, 1:148622272-148626199|+). We found that the upstream boundary of the seven other_circRNAs was located in a region of about 30 nucleotides around the start of exon5 (Figure 4A). The unique exonic circRNA of this region, 1:148622260-148626199|+ (Figure 4A), was the result of a backsplicing not supported by a canonical splicing signal. Indeed, we know that the motifs flanking canonical introns are recognized by the large spliceosome. These consensus motifs are ‘GT’ at the 5′ end of the intron and ‘AG’ at the 3′ end. The other_circRNA, (Figure 4A, 1:148622265-148626199|+) which was the most abundant circRNA from this region, may involve backsplicing supported by a canonical splicing signal (SS). The SS acceptor used in classical splicing does not seem to be the most “appropriate” to use for backsplicing. Therefore, there would be a competition to find another SS acceptor to perform backsplicing with the best efficiency. Nevertheless, one further other_circRNA (Figure 4A, 1:148622272-148626199|+) with a canonical splicing signal was observed, but with very low expression. Finding a canonical SS may not always be sufficient for selecting the backsplicing targets. The same kind of competition can take place to choose the branch point [36]. When we looked again at the eight circRNAs from *MORC3* that share their downstream boundary (end of exon7), we noted that they all were detected in the 63 total-RNAseq datasets, but not in any of the 63 mRNAseq datasets (Figure 4A). It can be assumed that they do exist *in-vivo*, but are created by imperfect backsplicing. We propose to call these circRNAs aberrant-exonic circRNAs, because they are generated by backsplicing, but to not fit into the exonic circRNA pattern. A second example of aberrant-exonic circRNAs is reported in Figure 4B and concerns the *LIFR* gene (ENSBTAG00000010423). In this case, the aberrant-exonic circRNA (20:35900693-35905466|+) is more lowly expressed than the exonic circRNA (20:35900693-35905467|+). Only the second is provided by backsplicing and supported by a canonical splicing signal. Observations on *MORC3* and *LIFR* exonic and aberrant-exonic circRNAs support the idea that backsplicing may be more demanding in terms of canonical SS than classical splicing [5; 28; 49; 50; 51].

**Figure 4:**
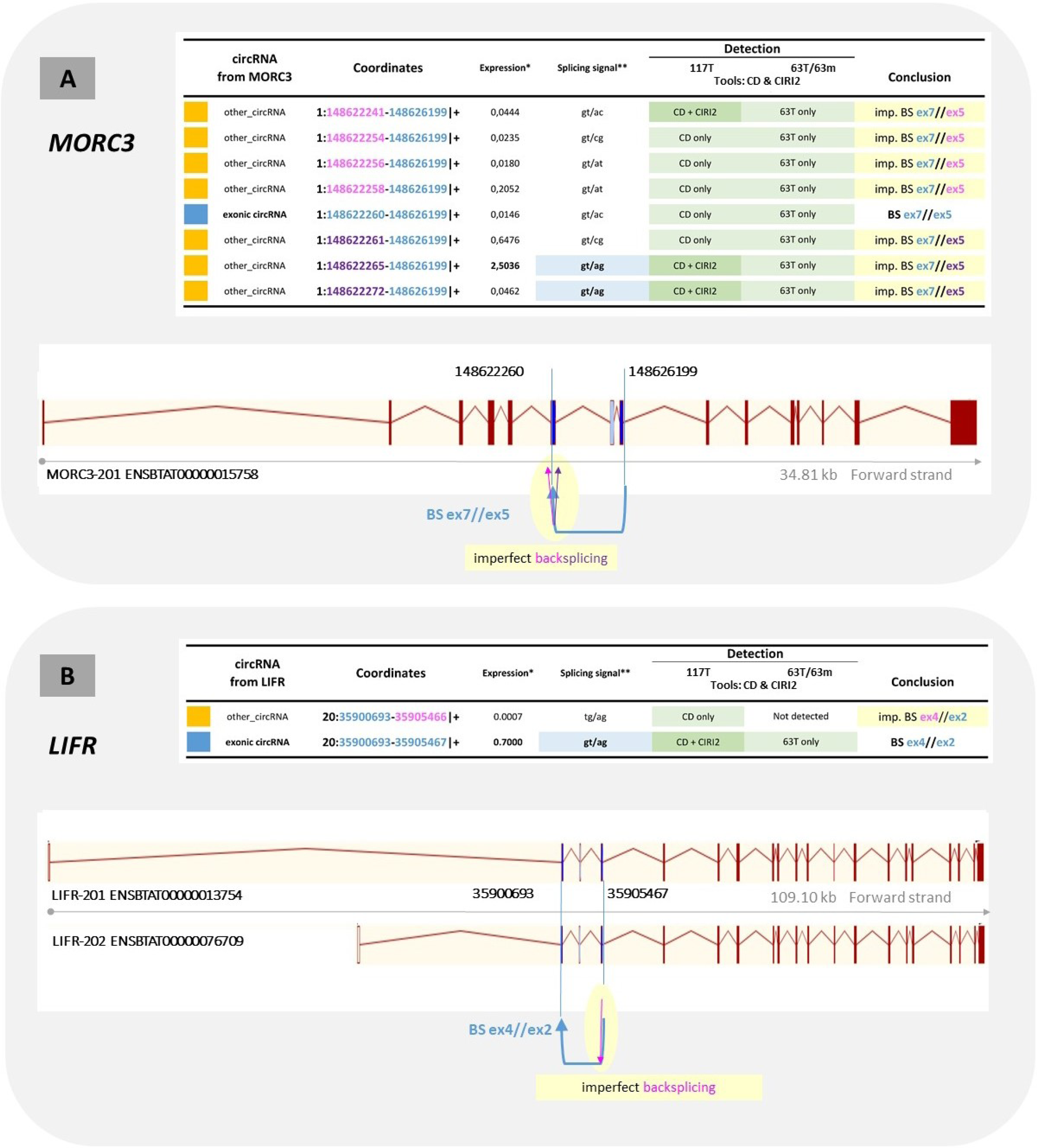
Perfect and imperfect backsplicing in two genomic regions. **(A)** In *MORC3* region (ENSBTAG0000000011876; BTA1: 148,604-148,640 Mb), CD identified 7 other_circRNAs and 1 exonic circRNAs. Three of the seven other circRNAs are also detected by CIRI2. None of the eight circRNAs from *MORC3* are detected in 63m by CD. **(B)** In *LIFR* region (ENSBTAG00000010423; BTA20: 35,840-35,950 Mb), CD identified 1 other_circRNA and 1 exonic circRNA. Only the exonic circRNA is also detected by CIRI2. These 7+1 other_circRNAs are probably generated by imperfect backsplicing * Expression: average of the expression observed for 117 tissue samples ** Canonical motifs of a backsplicing exon7-exon5 are localized in the 5’ boundary (gt) of the intron 7/8 and in the 3’boundary of the intron-4 ⁄ 5 (ag). Figure constructed from data and drawing uploaded on December 15, 2023 from the Ensembl bovine genome database (http://www.ensembl.org/Bos_taurus/Info/Index).

All examples of aberrant-exonic circRNAs shown in Figure 4 had a circRNA boundary that corresponded exactly to one exon boundary. Among the 36,215 CD-other_circRNAs, we detected 3,974 with this feature. Nevertheless, only 11.3% of them (450/3,974) have their second genomic coordinate in the region of the boundary of an exon (-30/+30 nt from the boundary). This statistic suggests that aberrant-exonic circRNAs are most likely very rare among other_circRNAs.

#### Parallel analyses performed in total-RNAseq and mRNAseq reveals artificial circRNAs

The availability of bovine paired datasets was a good opportunity to perform a circRNA detection in total-RNAseq and mRNAseq in parallel. We put together a new dataset of 63 samples from 11 tissues (see STab-1) with high-quality mRNA and total-RNA data available. Even though we avoided including samples from XL sequencing for total-RNAseq, the number of reads available for total-RNAseq was higher than for mRNAseq.

The 63 total-RNAseq dataset (63T, Figure 5A) identified over 35,000 circRNAs, while the 63 mRNAseq dataset (63m) identified 4,579 circRNAs (Figure 5B). The high number of circRNAs detected in 63m, which represents more than 10% of the number detected in 63T, was completely inconsistent with an expected background. Indeed, we expected to find circRNAs existing *in-vivo* but resulting from non-specific binding to oligo(dT) beads and *in-silico* artif_circRNAs. Moreover, these two possible types of circRNAs present in mRNAseq were expected in total-RNAseq. Upon examining the 4,579 circRNAs from 63m, it was observed that 63.4% (2,901 out of 4,579) had not been previously identified, i.e. in 117T (Figure 5B). Consequently, all circRNAs detected in mRNAseq were deemed unreliable and artificial. Additionally, it is important to determine the source(s) of these artif_circRNAs.

**Figure 5:**
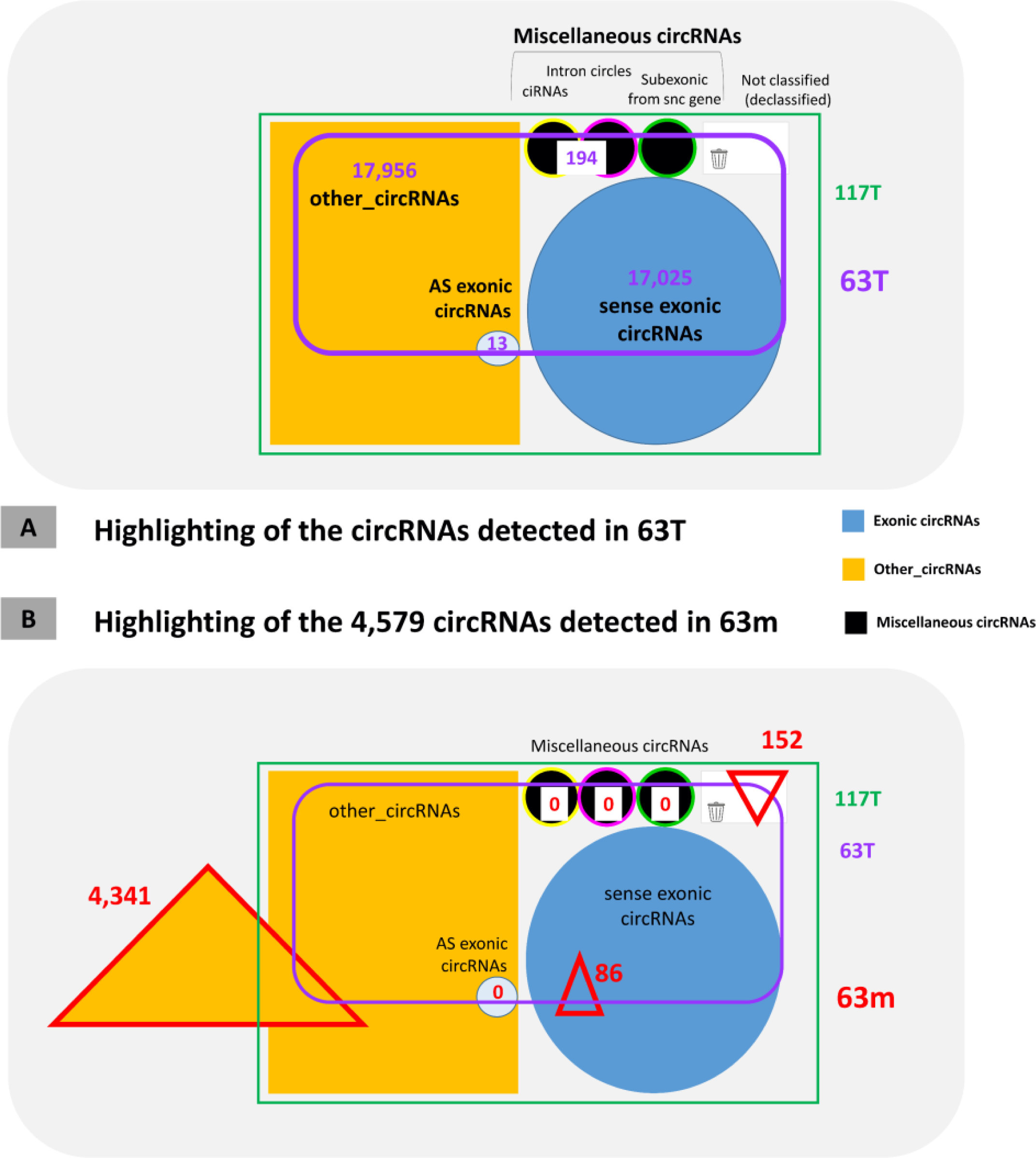
Analysis of circRNAs detected in mRNAseq. **(A)** Among the circRNAs detected in the 63 total-RNAseq dataset (63T, symbolized by the purple frame), we recognized 17,025 exonic circRNAs, 194 intronic circRNAs, and 17,956 other_circRNAs already identified in the 117T datasets. **(B)** In the 63 mRNAseq dataset (63m), 4,579 circRNAs were detected (they were represented by three red triangles), of which 2,901 (63.4%) had never been described before, i.e. identified in 118T. Neither miscellaneous circRNAs nor AS exonic circRNAs were detected in 63m (represented by three black discs and one light blue disc, respectively). Among the 4,341 other_circRNAs identified in 63m, 2,812 are novel. Among the 86 exonic circRNAs identified in 63m and already detected in 117T, 10 had not been detected in 63T.

In mRNAseq datasets (63m), we did not detect any intronic circles, ciRNAs, or sub-exonic circRNAs from snc genes. In addition, no AS-exonic circRNA was detected in the 63m (13 and 20 were detected in 63T and 117T, respectively) and the eight aberrant-exonic circRNAs reported in Table 3 and detected in 63T were also absent in the 63m (Figure 5B). We have no reason to suspect these different types of circRNAs as unreliable.

In the 63m dataset, we identified 86 sense exonic circRNAs from the list of circRNAs found in 117T (Figure 5B). The level of artif_circRNAs is very, very low among exonic circRNAs. We cannot be as assertive among intronic circRNAs or sub-exonic from snc, as the samples were much poorer.

In the 63m dataset, we retained 4,341 circRNAs as other_circRNAs but 2,812 (64.8%) have never been detected in total-RNAseq. We detected a higher proportion of other_circRNAs with a small genomic size (70-159 nt) in 63m than in 63T (data reported in Res_Adoc-2 showed a statistically significant difference). This observation is certainly related to the higher proportion of other_circRNA that we could have classified as sub-exonic from multi-exonic genes (both genomic coordinates are located in the same exon, nuclear chromosomes and mitochondrial genome, Ensembl annotation, see Res_Adoc-6A). The fact that we observed the same number of these sub-exonic circRNAs per million uniquely mapped reads in 63m and 63T (Res_Adoc-6B) does not support their reliability, as their genesis seems to be automatic or mechanical.

In an attempt to get a clearer picture of the reliability of other_circRNAs, we proposed to focus on three regions (see also Res_Adoc-7). In the *albumin* gene region, sample 63T is as informative as 117T for other_circRNAs. Most of the other_circRNAs identified in this region (57/59) have both genomic coordinates in exonic sequences (in the same exon (sub-exonic) or in two exons). Since the five new other_circRNAs detected in 63m have the same feature (5/5), it was difficult to conclude that the 32 other_circRNAs identified only by total-RNAseq were reliable. The analysis of the other_circRNAs from the mitochondrial genome detected in 63m led us to consider them all unreliable (see Res_Adoc-7). This is not surprising because the characteristics of other_circRNAs from the mitochondrial genome are very close to those of sub-exonic circRNAs from multi-exonic genes. In contrast to, the statistics about the other_circRNAs detected in the *Defensin* gene region (Res_Adoc-7) suggest that a very large proportion of them are reliable. Undeniably, this 63m/63T comparative study casts doubt on the reliability of at least a large proportion of the other_circRNAs.

#### Complementary analyses performed with CIRI2

Using CIRI2 [24] on the 117 total-RNAseq datasets, 58,373 CIRI-circRNAs were detected with at least two reads spanning the circular junction. Even though we consider this too low a threshold, it is the upper limit for initial analysis implemented in the CIRI2 program. For example, 350 CIRI-circRNAs were identified in the *Defensin* region. Among the 23,926 exonic circRNAs identified by CD, 20,531 were also identified as CIRI-circRNAs (Figure 6A). The overall confirmation rate for CD-exonic circRNAs is 85.8%, but only 2/27 for exonic circRNAs from the *Defensin* region. We also identified 2,305 other-circRNAs identified by CD among the CIRI-circRNAs (Figure 6A). The confirmation rate for CD-other_circRNAs is 6.4% (only 17/3,160 for other_circRNAs from the *Defensin* region). When the annotation of these 58,373 CIRI-circRNAs was performed with CD, 48,310 putative exonic circRNAs were suggested. Among the 18 putative miscellaneous circRNAs, we retained three ciRNAs, nine intron circles 3 sub-exonic circRNAs from snc genes (Figure 6B). These data further defined a set of 10,102 CIRI-other_circRNAs (Figure 6B). Of these 10,081 circRNAs, only 111 (1.1%) had a small genomic size (135-160 nt) (Res_Adoc-2). Furthermore, none of them was from the mitochondrial genome. When we continued the comparisons of the features of these CIRI-other_circRNAs (see for details Res_Adoc-3), we were able to conclude that the other_circRNAs identified by CIRI2 were not the same as those identified by CD. These observations are not very surprising as the design of these two bioinformatic tools is different.

**Figure 6:**
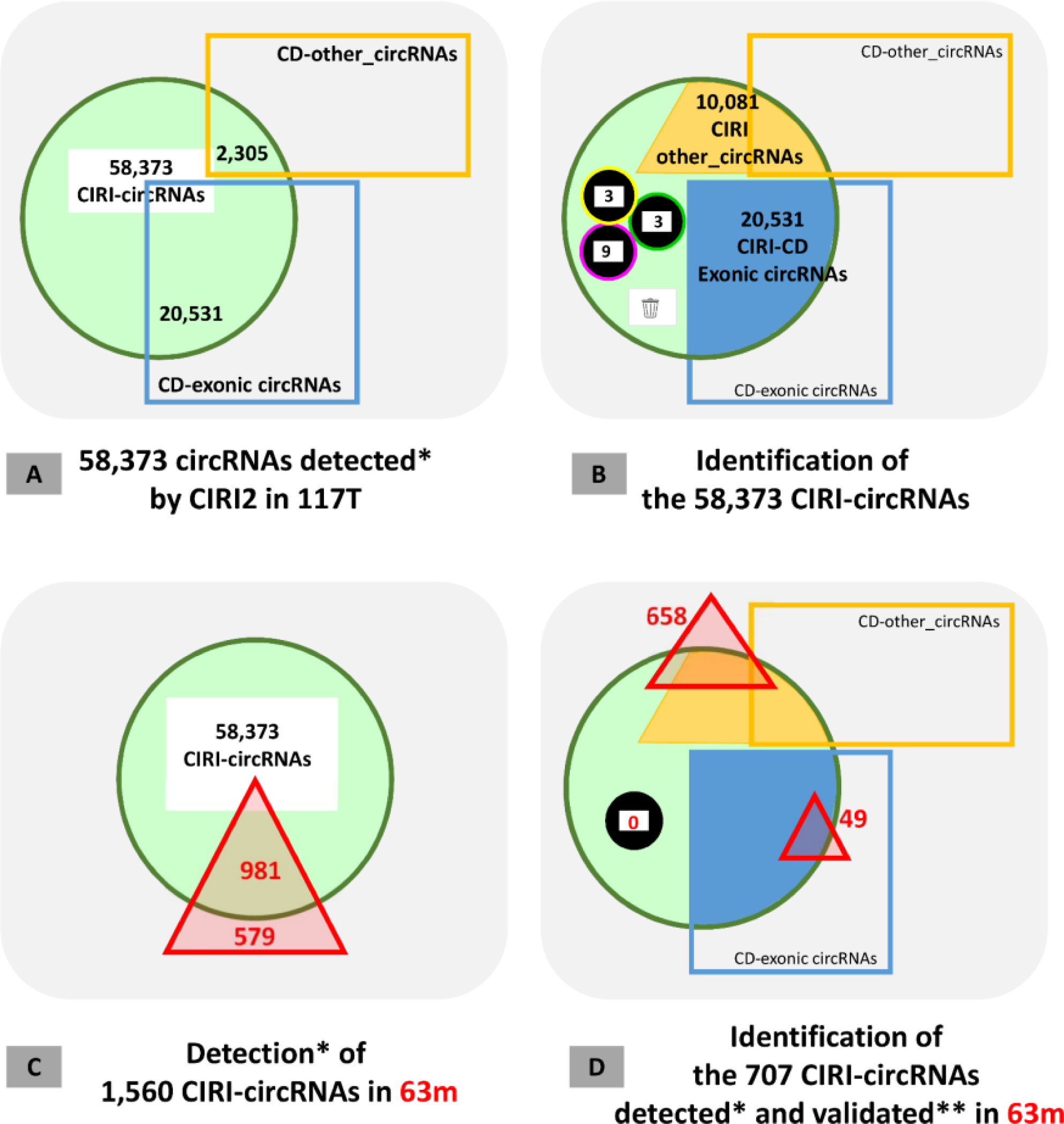
Analysis of circRNAs detected by CIRI2. **(A)** CircRNAs detected by CIRI2 in 117T were represented by a green circle. Those have already been detected and annotated by CD were highlighted by a bleu rectangle (CD-exonic circRNAs) and by an orange rectangle (CD-other_circRNAs). **(B)** Among the CIRI-circRNAs from 117T, we identified 20,531 exonic circRNAs already identified by CD, 15 miscellaneous circRNAs (represented by three black discs) and 10,081 CIRI-other_circRNAs. **(C)** 1,560 CIRI-circRNAs were detected* in 63m (represented by a red triangle) **(D)** 707 CIRI-circRNAs were detected* and validated** in 63m (represented by two red triangles corresponding to exonic circRNA and other-circRNAs respectively). No miscellaneous circRNAs detected in 63m (represented by one black disc). * CIRI2 retained a circRNA “as detected” when at least two reads spanning the circular junction in at least one individual dataset. ** for the circRNAs detected in 63m by CIRI2, we considered as “validated” only those detected by at least five reads spanning the circular junction in at least one individual dataset.

Subsequently, we performed a new detection of circRNAs in the 63m dataset using CIRI2. Out of the 1,560 CIRI-circRNAs detected, 579 were not present in the list of 58,373 CIRI-circRNAs detected in 117T (Figure 6C). The detection in 63m of these previously undescribed CIRI-circRNAs (37.1%) confirmed that all circRNAs detected in mRNAseq can be considered artificial. However, we only kept the 707 CIRI-circRNAs that were detected with at least five reads spanning the circular junction, which were validated as artificial circRNAs. Among them, we recognized 49 CD-CIRI exonic circRNAs from the list established on 117T by CD. We can conclude that the list of CIRI-circRNAs detected in mRNAseq included no more than 4% of the validated exonic circRNAs. These 49 circRNAs which were previously annotated as exonic circRNAs are now considered as artif_circRNAs, since they were detected in mRNAseq. Similar to CD, CIRI2 can also identify 63 circRNAs that were not detected in 63T. For example, 6 out of 49 CD-CIRI exonic circRNAs were identified.

### Refinement of the list of exonic circRNAs

#### Identification of 103 artificial circRNAs among the list of exonic circRNAs

We identified 86 and 49 exonic circRNAs as artificial circRNAs from the analyses of 63T/63m by CD and CIRI2, respectively. Since 32 were identified by the two approaches (see STab-5), we suggest that 103 circRNAs previously annotated as exonic circRNAs are artif_circRNAs.

When we examined backsplicing falsely identified at the origin of these 103 artif_circRNAs, we found 2/5 from two Ensembl exons (42), 2/5 from two MSTRG exons (40) and 1/5 from mixed pairs (21). These observations showed a statistically significant difference with the observations made on the list of 23,926 exonic circRNAs where 77% of backsplicing involved a pair of Ensembl exons. (Res_Adoc-1, chisq_test with p-value <2.2 10^-16^).

Among this list of 103 artif_circRNAs, we find the circRNA with the highest average expression across all of the 117 samples (2:18153915-18180018|+). This circRNA was actually only detected in the two neonatal animals, which were also the two Belgian animals. It could be an artif_circRNA generated by differences in this genomic region (*TTN*), specific for these two animals of the same genetic origin (Holstein Friesian). Nevertheless, it is surprising that CD and CIRI2 detected it in 117T, while only CIRI2 detected it in 63m. Among the set of 103 artif_circRNAs, we also noticed the presence of a cluster of 11 circRNAs from a region on BTA23 (28.52-28.72 Mb) containing a part of the major histocompatibility complex (MHC) class I genes. These 11 circRNAs were ‘linked’ by exon(s) identified as involved in (false) backsplicing and originating from the same MSTRG gene. This cluster included the circRNA with the highest average expression across the 117 samples, but was in fact only expressed in the tissues of neonatal animals. In the same region, 63 other_circRNAs were characterized in 63T, but 54 were invalidated by the analysis of 63m. Moreover, 26 novel other_circRNAs were detected in 63m.

#### Fine annotation of exonic circRNAs reveals some artificial annotations

Using two separate exon lists (Left_exons and Right_exons), a second annotation called minimal_annotation was performed for each of the 23,926 exonic circRNAs. In this way, we identified the two exons involved in each backsplicing, and when alternatives were possible, only the smaller exon was considered, regardless of the name of the parental gene (for details see the Materials and Methods section and M&M_Adoc-5). This minimal_annotation led to the characterization of a larger fraction of exonic circRNAs annotated with an Ensembl exon and an MSTRG exon compared to the classical CD-based circRNA annotation (Res_Adoc-1). Only 30,831 different exons (4.8%) out of the 683,396 described exons of the bovine genome (it would be more correct to take into account only the 636,307 considered for the minimal_annotation) were involved in the generation of exonic circRNAs.

We can describe the group of bovine exons involved in backsplicing by a mean size of 188 bp and a median size of 133 bp. These exons appear to be larger than those characterized as involved in exonic circRNAs in human HEK cells (160-165 bp for the mean size, [47]) or in exonic circRNAs from porcine testis (148 bp for the mean size, [17]). We detected 48 circRNAs annotated with two overlapping exons among the 23,926 exonic circRNAs, however, these two exons cannot be associated in the same transcript. Thus, these 48 circRNAs are not true *in-vivo* exonic circRNAs. Among them, we found 23 of the 27 circRNAs identified as exonic circRNAs from the *Defensin* gene. Our analysis also led to the identification of 1,025 single exon circRNAs (see the M&M_Adoc-4). The average length of these exons is 605 bp. This size is consistent with the one observed for a single-exon circRNAs from porcine testis (647 nt, [17]) or human HEK cells (709 nt, [47]). Among the list of 1,025 single exon circRNAs, we noted that seven were originating from the same parental gene (ENSBTAG00000006907, *Nebulin*, *NEB*), which is in itself suspicious. Moreover, we noted that four of them were detected in 63m by CD, and, thus, were already suspected to be artif_circRNAs. An eighth exonic circRNAs from the same region seemed suspicious, since it involved a BS between two of the same exons. We suggested not retaining these eight circRNAs from the *Nebulin* region as exonic circRNA. Since single exon circRNAs range in size from 76 to 6,723 nt, we can suspect that exons larger than 7 kb are likely too large to be involved in backsplicing. When the list of 23,926 exonic circRNAs was examined in regard of the size of both exons involved in the backsplicing, we decided to not retain as exonic circRNA nine circRNAs involving at least one very large (15-38 kb) exon. All are MSTRG exons from the BovReg annotation.

This deep exon-based annotation allowed the identification of 48 (overlapping exons) + 8 circRNAs (many single exon circRNAs from the same gene) + 9 (very large exon involved), i.e. 65 circRNAs that were initially described as exonic circRNAs but share an annotation casting doubt on their true *in-vivo* existence.

#### Discovery of 26 exonic circRNAs that were very suspicious

When the list of 23,926 exonic circRNAs was examined with respect to the size of the genomic region defined by their two genomic coordinates, we found 26 circRNAs that defined a region of up to 500 kb (0.1%). Among them, we did not find any exonic circRNAs identified as the result of backsplicing between two Ensembl exons. The first with this feature defines a region of 483 kb. In addition, in CD-other_circRNAs we identified 380 circRNAs with this feature (1%) and CIRI2 considers only circRNAs defining a genomic interval <200 kb. We considered these 26 circRNAs previously annotated as exonic circRNAs too suspicious to be reliable circRNAs, the probability that they are artif_circRNAs is very high.

#### The list of exonic circRNAs included 189 false entries

In addition to the 103 artif_circRNAs highlighted the analyses of 63T/63m by CD and CIRI2, and to the 26 probable artif_circRNAs highlighted by the examination of the size of the genomic interval defining the circRNA, the process of fine annotation led to the highlighting of 65 artificial annotations. As a result (see also SDoc-6), the list of exonic circRNAs was purified from 189 units and only 23,737 exonic circRNAs were considered for further analyses. The list of the 189 discarded exonic circRNAs is provided in STab-5.

### Bovine circular transcriptome

For these analyses, we first considered the 117 samples and then a group of 15 tissues for which samples were available from the two young animals and at least three juvenile or old animals. We detected an average of 5,329 exonic circRNAs with non-null expression in each of the 117 samples (Figure 7A), but only 1,711 exonic circRNAs (Figure 7B) with a notable expression. When we looked at the individual sample scale, “the number of exonic circRNAs with non-null expression” (Figure 7A) showed less homogeneity per tissue than “the number of exonic circRNAs with notable expression” (Figure 7B). Considering these two criteria for the number of expressed circRNAs, we observed a similar ranking for the 5 or 6 animals in only two tissues out of 15 (cerebellum and spleen) (Figure 7A, 7B). Regarding the testis with two samples, we noted that the numbers of expressed circular exonic RNAs evaluated by the two criteria (Figure 7A, 7B) are concordant and conclude that this number decreases with age in bovine. These results are consistent with our previous work characterizing circular exonic RNAs in tissues from three livestock species [27]. The cerebellum sample from the juvenile female showed the highest mean values for these two criteria (Figure 7A, 7B).

**Figure 7:**
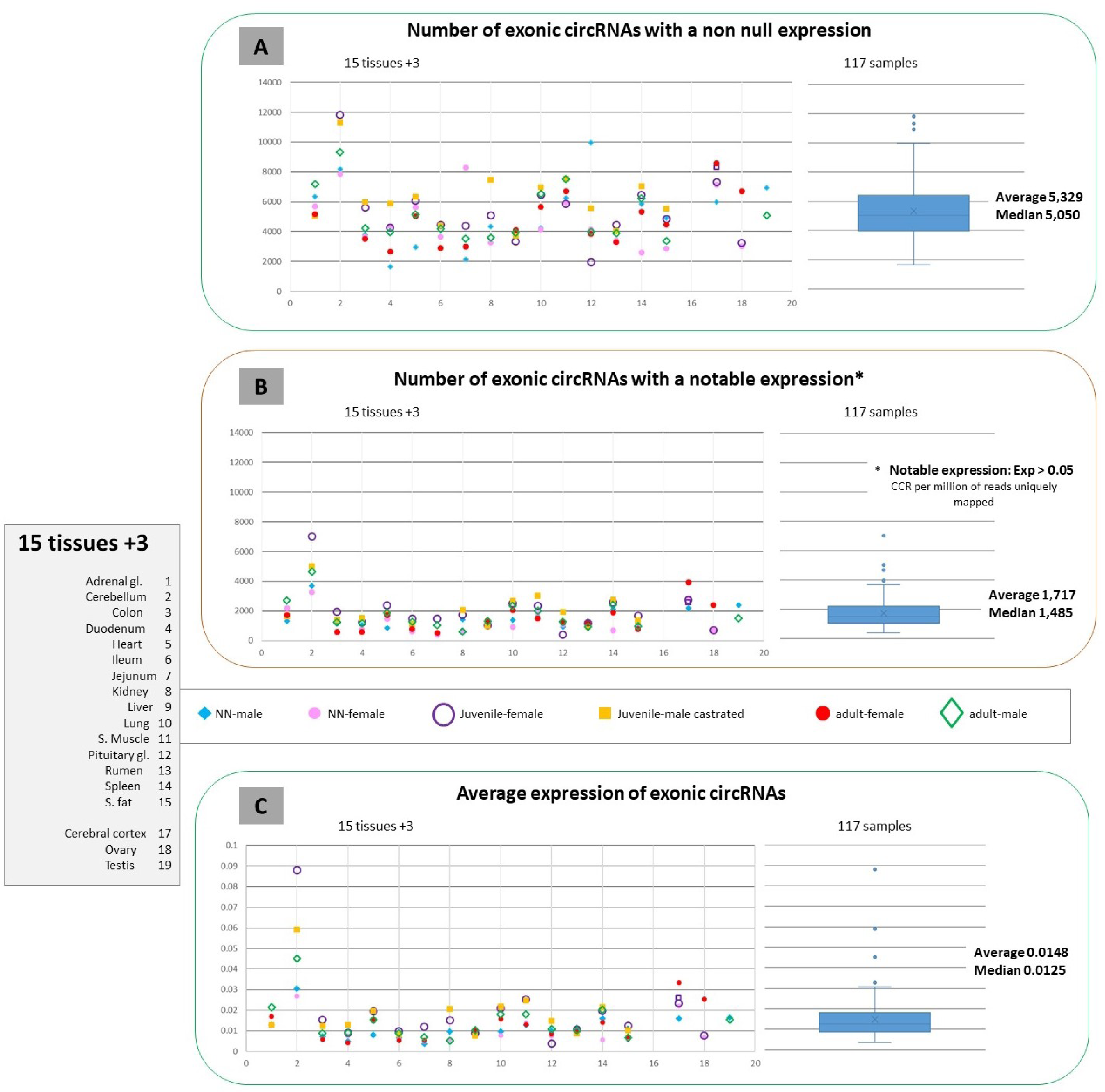
Analyses of the possible presence of 23,737 exonic circRNAs in bovine tissues/samples. All available samples for 15+3 tissues were considered in the left part and the 117 samples for the box plot shown in the right part. **(A)** and **(B)** represent a number of exonic circRNAs per million of reads uniquely mapped. **(C)** is dedicated to the observed expression, which is a number of CCRs per million of reads uniquely mapped. We defined “notable expression” as expression above 0.05. To make these 3 diagrams easier to read, they are also available in large format in Res_Adoc-8. The three tissue samples from neonate animals (jejunum-female, rumen-male, pancreas-male) that were sequenced at great depth are indistinguishable from the others.

For each tissue sample, the average expression level across each of the 23,737 exonic circRNAs (Figure 7C) was calculated. Among the four samples showing the highest expression level of the 117 samples, three were from the cerebellum (Figure 7C). The cerebellum differs from the other 14 tissues by the highest mean expression values per tissue (Figure 7C). The cerebellum was also distinguished by the variability in expression level that exists between samples (Figure 7C).

For the three criteria considered (Figure 7A, 7B, 7C), we observed that the XL sequencing, applied to three samples, did not affect the results. For these three criteria, the cerebellum from the juvenile female presented always the highest expression levels. This sample is also undoubtedly the sample with the most diverse circular transcriptome among the 117 samples considered here (Figure 8A, 8B1). In contrast, the circular transcriptome can be described as poor in terms of diversity, complexity and expression level for digestive tissues. One of the ‘poorest’ circular transcriptomes considered here is that of the jejunum from the neonatal male (Figure 8B2). Intermediate to the two extreme transcriptomes for cerebellum (rich) and jejunum (poor), are e.g. the testis and the ovary of adult animals (Figure 8B3 and 8B4).

**Figure 8:**
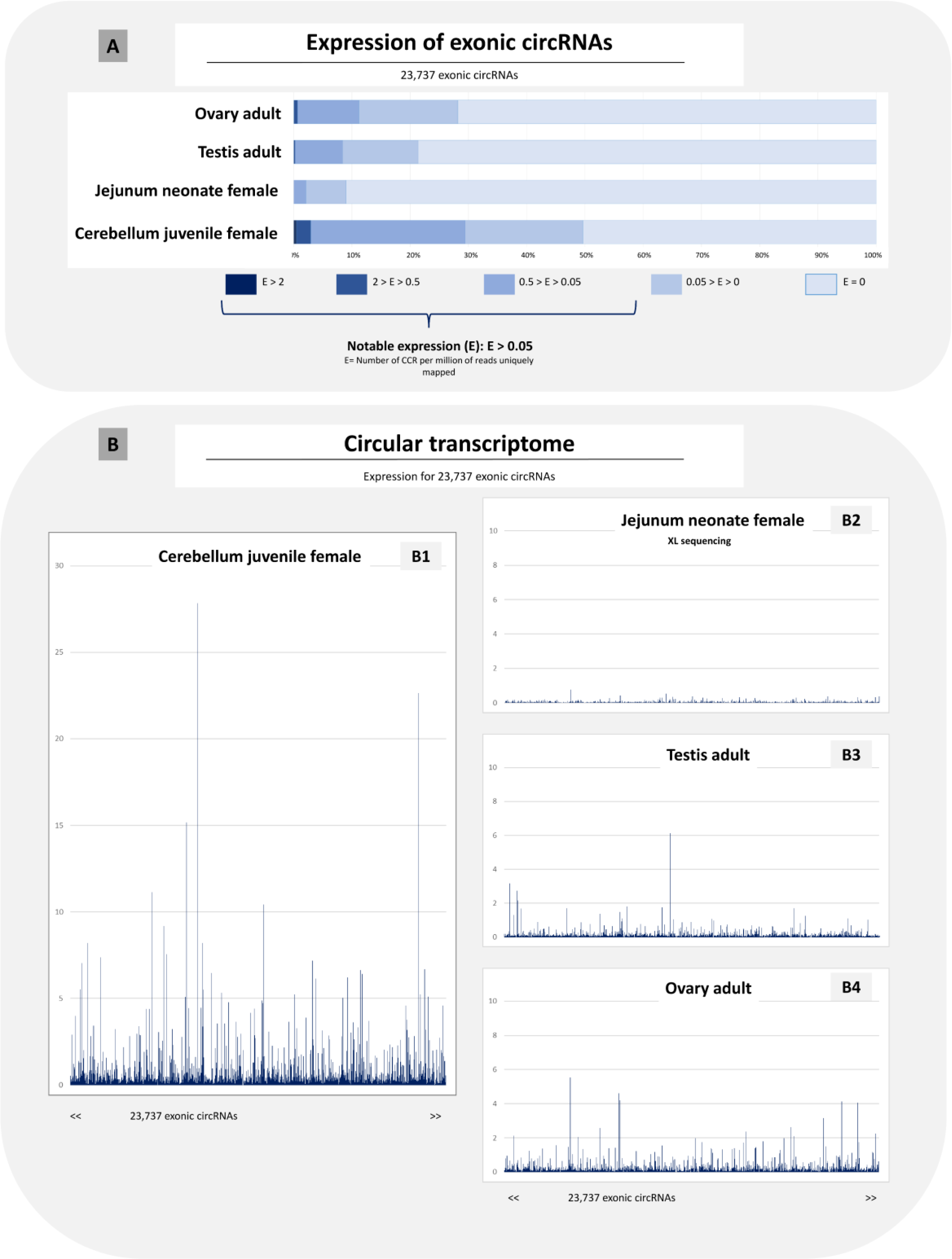
Expression analysis of 23,737 exonic circRNAs in four samples. The expression of a circRNA is defined as the number of CCRs per million of reads uniquely mapped. **(A)** Transcriptome composition and comparison of the four samples. **(B)** Schematic representation of four individual transcriptomes at the same scale. Other analyses concerning the jejunum neonate female and the adult testis are shown in Figure 2C.

#### To highlight the tissue specificity of the circular transcriptome by using a small panel of exonic circRNAs

The circular transcriptome is very complex and as such, it was important to determine if it is possible to reduce the complexity to better identify tissue specificities. We analyzed the tissue specificity of the circular transcriptome by considering a panel of exonic circRNAs. To avoid evaluating a tissue by a single dataset, we selected 15 tissues where samples were available for the two youngest and at least three of the oldest animals. In addition, we considered the five tissues where samples were available for the two young animals. To this end, we performed hierarchical cluster analysis (HCA). The ideal result would be to find a clustering of the circular transcriptomes by tissues. Three HCAs were performed, with (1) 96 samples (15 tissues with 5/6 animals + 5 tissues with only young animals), (2) 56 samples (15 tissues with only juvenile or old animals), (3) 40 samples (20 tissues with only young animals). Although we explored 23,737 exonic circRNAs in 117 samples, only 386 to 6,995 had a notable expression in a given sample, and three samples were sequenced at a higher depth with XL sequencing. We wanted to prevent circRNAs with very low expression from becoming the discriminators. To construct an exonic circRNA tissue evaluation panel we included in the respective list of circRNA those samples which were the top-150 exonic circRNAs ranked according to their expression level in any of the 116 samples (we did not include circRNAs data obtained from the second total-RNAseq from the cerebral cortex of the juvenile castrated male). This method resulted in a panel of 1,749 exonic circRNAs (list available in STab-6).

We began the HCA by considering the expression of 1,749 exonic circRNAs in 96 samples. With normalization performed using the log-binary method (Figure 9), we observed tissue-wise clustering of all samples for nine tissues within the group of 15 tissues (cerebellum, muscle, heart, kidney, adrenal gland, lung, spleen, liver, and rumen). When we considered only the oldest animals, we noted clustering of two additional tissues where the two youngest animals were still available (fat and pituitary gland). In addition, the two samples from the youngest animals clustered together for thyroid, pancreas and cerebral cortex (tissues where samples from the oldest animals were not available). As the age of animals had an effect on the clustering pattern, we proposed to analyze separately young and juvenile/old animals. The clustering using the log-binary method considering only the 56 samples from the oldest animals and only 40 samples from the youngest animals were consistent with results observed on HCAs built with 96 samples (Res_Adoc-9A).

**Figure 9:**
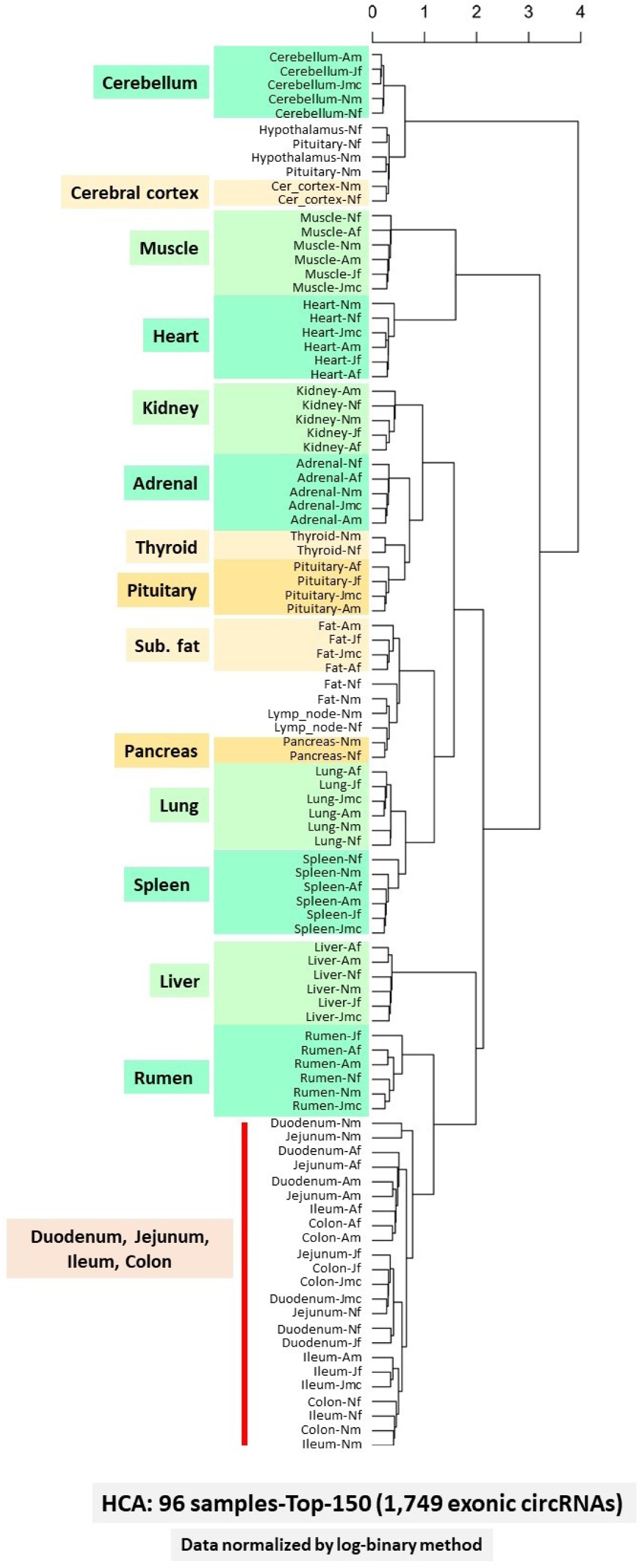
Hierarchical clustering analysis (HCA). HCA built using the “ward” agglomeration method and Pearson correlations as distance on the expression of 1,749 exonic circRNAs (panel top-150, composition in STab-6) in 96 samples. Each sample was labeled with a name composed as “tissue-agesex” where age=N (neonate) or J (juvenile) or A (adult). When the clustering corresponds exactly at the expected (by tissue) the corresponding tissue was underlined in green (5 or 6 animals) or in yellow (2 or 4 animals).

Several panels increasing or decreasing the number of top exonic circRNAs ranked according to their expression were created with the expectation of improving the clustering (increasing the number of tissues where all samples clustered according to tissue). HCAs (96, 56, and 40 samples) were constructed with data normalized by the log-binary and standard score methods. Differences from the respective reference results (HCA obtained with top-150) were observed, but they were mainly negative differences (see Res_Adoc-9). No clear improvement was observed regardless of the normalization method used. These analyses were inconclusive for the four digestive tissues (duodenum, jejunum, ileum, and colon), which did not show a tissue- or organ-specific clustering pattern. In addition, we often observed a degradation of the clustering quality, especially when clustering small groups of samples (56 and 40).

These analyses showed that the top-100 and -150 panels are the most efficient, whatever the normalization method used, and even we considered only a subset of the 116 initial samples. The top-150 with 1,749 exonic circRNAs (7.4% of reliable exonic circRNAs) can be considered as a reference. We emphasized that this panel included the most highly expressed exonic circRNAs (top-150 for each of 116 samples). The lists of exonic circRNAs constituting the top-100 to top-250 panels are available in STab-6.

#### Analysis of reproductive tissues

To analyze the circular transcriptome of reproductive tissues, we performed a PCA on the expression of circRNAs from the reference panel (1,749 exonic circRNAs, panel top-150) in these tissues (uterus, uterine horn, testis, and ovary). In addition, we considered the adrenal and pituitary gland samples. Initially, we considered these 6 tissues and 19 individual samples in total (Figure 10A). The first two and the first four PC dimensions explained 42.00 and 66.64% of the variance, respectively. The first dimensions allowed us to separate the samples from the pituitary gland into two groups (Figure 10A1). We found that these groups did not reflect the age or the sex of the animals sampled. The most interesting element was probably that the testis of the adult animal appeared as an outlier in the dim-3 (Figure 10A2). Since we were not convinced that the consideration of the pituitary gland was informative, a second PCA was performed with 5 tissues and 13 individual samples (Figure 10B). The performance of this PCA was better than the previous one, as the first two and first four dimensions explained 54.11 and 71.51% of the variance, respectively. The dimension-1 allowed the individualization of the sample from adult testis (Tes-A on Figure 10B1). The first dimensions allowed us to separate the samples into two groups and two individual samples (the two testis) (Figure 10B1). The first group included all female reproductive tissues (uterus, uterus horn, and ovary). The second group included all samples from the adrenal gland. The dimensions-3 and -4 showed a proximity between the testis of the adult animal and the adrenal gland samples of both adult animals (male and female) (Figure 10B2).

**Figure 10:**
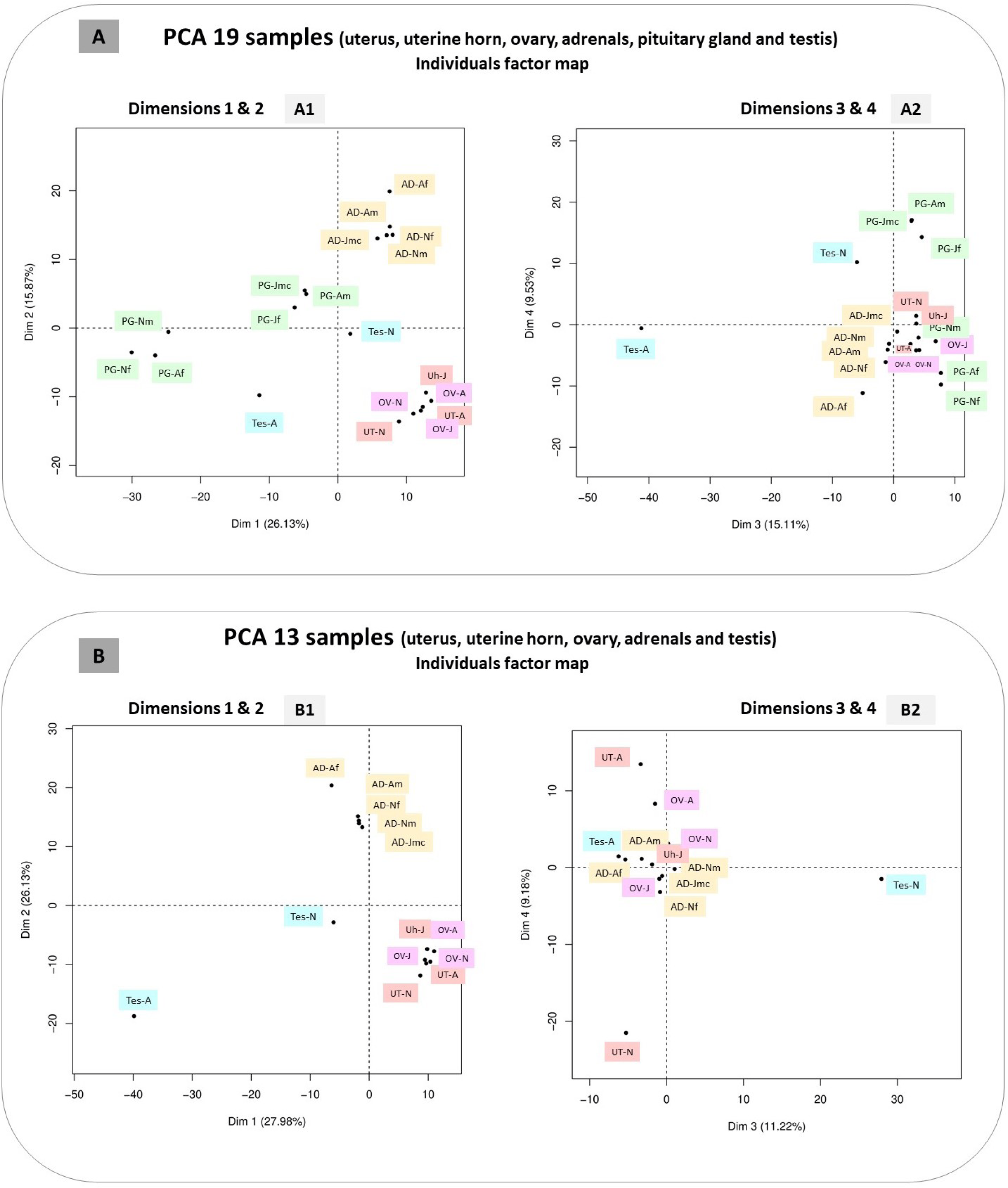
Principal component analyses (PCA). Both PCA were built on the expression of 1,749 exonic circRNAs (panel top-150). The plots show the individual factor maps, dimensions 1 and 2 on the left and dimensions 3 and 4 on the right. The readability of the labels on these plots has been manually improved. Samples from neonates were labeled -N, and -Nm- or -Nf when sex precision was useful. Samples from juveniles were symbolized by -J, and by -Jmc- or -Jf when the precision of the sex is useful (castrated male and female). Samples from adults were denoted -A, and - Am- or -Af when sex precision is useful**. (A)** Six tissues were considered: uterus (UT), uterine horn (Uh), ovary (OV), adrenal gland (AD), pituitary gland (PG), and testis (Tes). (**B)** Only samples from five tissues were considered (the six samples from PG were removed).

## Discussion

In this study, we performed a circRNA characterization in bovine tissues with total-RNAseq data generated in a standardized manner for the BovReg project. We avoided the agglomeration of other available datasets to minimize batch effects that make interpretation of results difficult [27]. This is because composition of the circRNA catalog depends not only on the sample considered but also on the tissue collection and preservation method, RNA isolation and sequencing library preparation protocols, and data analysis pipeline. In this study we detected 81% (7,071/8,723) of the exonic circRNAs that were previously detected by our previous work in a smaller subset of tissues in 2021 (muscle/liver/testis and detection by CircDetector [27]). Furthermore, when only exonic circRNAs that were identified in liver or muscle samples, are consider the detected percentage in this study relative to our previous study rose to 88% (3,565/4,050). Moreover, we observed that 86% of the exonic circRNAs identified and validated by CD were also detected by CIRI2. What was more surprising than the number or list of exonic circRNAs, was the low proportion of circRNAs that can be annotated as exonic circRNAs [27]. The low percentage of exonic circRNAs (40%) observed in this study was far lower than the value we observed in pigs [27; 28], though this value seems to vary depending on the tissue or the origin of the datasets [27]. The use of the new transcriptome annotation allowed a 191% increase in the number of identified exons, but did not allow a clear improvement in the percentage of circRNAs annotated as exonic circRNAs compared to our previous study [27]. However, of these exonic circRNAs, there are very few that have only been validated in one dataset, and as such they are likely to be more ‘reliable’ than the other_circRNAs where the majority are only validated in one dataset. The diversity in the population of other_circRNAs and the number of datasets considered led to a very low average percentage of exonic circRNAs. The exonic circRNAs characterized here are likely to be reliable circRNAs, based on the criteria defined by Chuang et al. in 2023 [52] while the other_circRNAs may not be.

The analyses conducted here showed that there were signals of circularization events in the reads obtained from mRNAseq, but that these were often never observed in total-RNAseq reads (CD and CIRI2 analyses). Moreover, these circRNAs are very rarely exonic circRNAs, even with CIRI2. This shows that Lv et al. [30] had not worked on mRNAseq data as they reported. The essential feature of the artificial circularization events detected in mRNAseq seems to be that they are not reproducible. We were somewhat surprised that not all artif_circRNAs belonged to the other_circRNA category, since at least 103 artif_circRNAs were detected among those annotated as exonic circRNAs. Conversely, the other_circRNA category did not contain only artif_circRNAs. Among the ‘reliable’ other_circRNAs, we identified a small number of aberrant exonic circRNAs and most of the circRNAs were found in the *Defensin* genomic region.

As previously described [27], clusters of other_circRNAs were characterized in several genomic regions known to be incompletely sequenced and incompletely assembled. We hypothesized that the presence of inverted regions in the assembled genome led to the mapping of some reads as artificial CCRs and to the identification of *in-silico* artif_circRNAs. We could have the same consequences in regions with gene clusters with segments of high homology, creating opportunities for misalignment. Therefore, we were not surprised to highlight more artif_circRNAs than real circRNAs in the MHC region. At the beginning of this study, we also thought that the *Defensin* region would be a good example of a region producing *in-silico* artif_circRNAs [27]. Analysis of the other_circRNAs present in 63T and 63m clearly showed that the other_circRNAs identified in the *Defensin* region seem “reliable” circRNAs. It is possible that their identification as exonic circRNAs failed due to small gaps or errors at the boundaries of the exons. However, the statistics do not support this simple explanation. We showed that this region is capable of producing circRNAs (over 3,000), which seem reliable because they were not detected in mRNAseq data, but only four have been identified as exonic circRNAs. This region is particularly difficult to understand, undoubtedly due to a mixture of problems (e.g., sequencing/assembling, highly homologous genes with copy number variations between individuals, and non-poly(A) transcripts).

The consideration of mRNAseq in addition of total-RNAseq led to the identification of 103+26 artificial circRNAs among the list of exonic circRNAs. We can propose several hypotheses to explain these backsplicing falsely identified (Figure 11): (1) the existence of inverted genomic sequences in the assembly. (2) The existence of genomic sequences with high similarity in the reference genome (gene family organized in clusters). (3) The presence of small regions in the genome of the affected animals with inverted genomic sequences or with chromosomal rearrangements. (4) Confusion with the identification of transcripts resulting from trans-splicing. (5) Possible template switching during reverse transcription in the library preparation process. The first three could be assimilated to *in-silico* circularization, and the fourth is due to an *in-vivo* event, but it is not a circularization event [13; 29]. The third hypothesis may be illustrated by the artif_circRNA (2:18153915-18180018|+) detected in the *TTN* region, but only in the two Belgian animals. It is difficult to imagine that the hypothetical fifth event (*in-vitro*) would reproducibly lead to a circular junction identifiable as an exonic circRNA. We noted that the consideration of exons novelty annotated by BovReg increased the risk to annotate an (artificial) circRNA as an exonic circRNA. In addition, we were aware to take a risk by accepting circRNAs with backsplicing between a mixed pair of exons (Ensembl/MSTRG).

**Figure 11:**
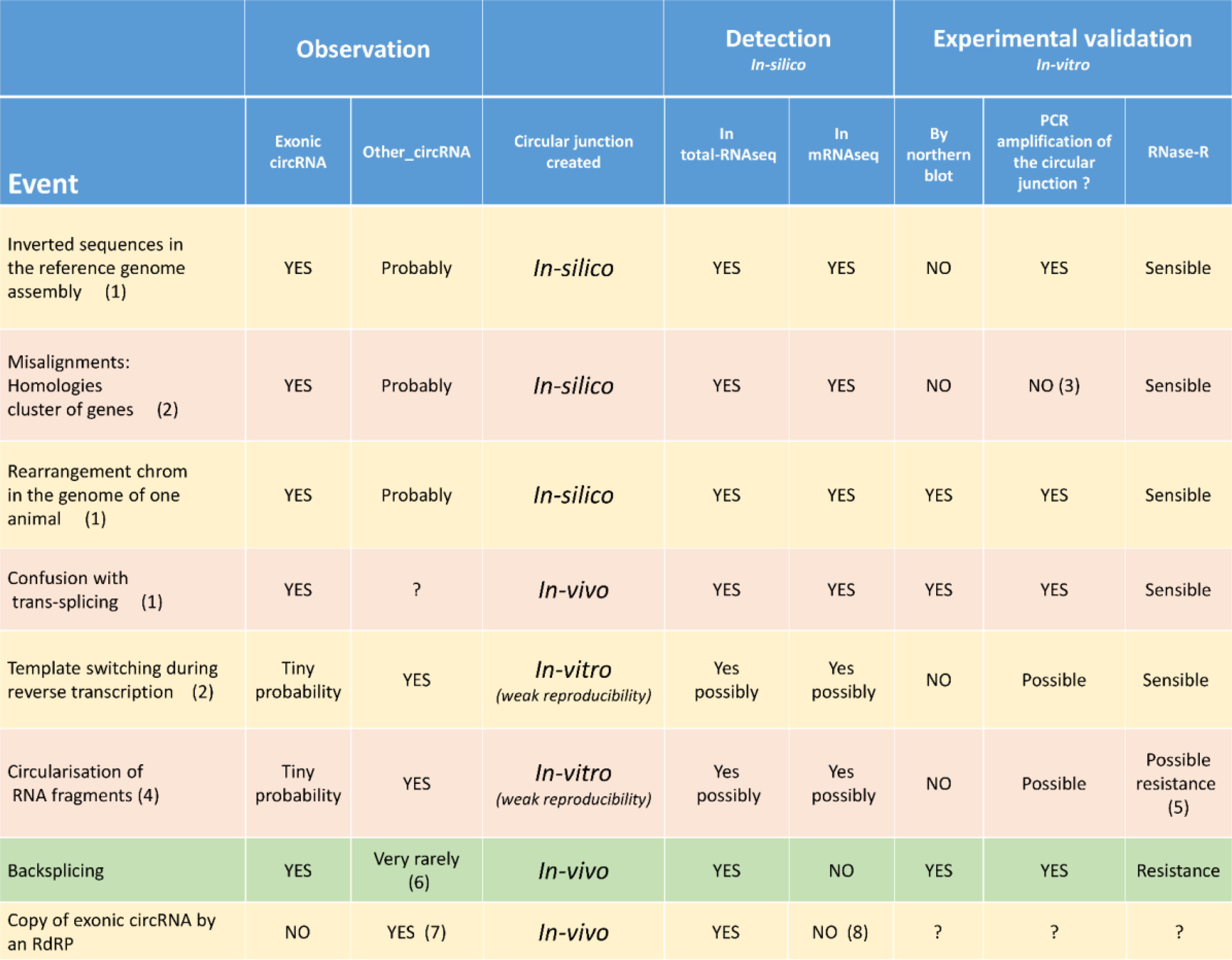
List and characteristics of different events leading to the formation of a circular junction. Six (hypothetical) events leading to the identification of artificial circRNA are listed on an orange or yellow background. Backsplicing leading to exonic circRNA is described on a green background. Additional information: (1) The transcript containing the “circular junction” exists but is not circular. (2) The transcript containing the circular junction is not present. (3) The cDNA containing the circular junction is not present. (4) The transcript containing the circular junction is present and is circular. (5) In addition, the junction may have been created after RNase-R action. (6) Aberrant exonic circRNAs obtained after imperfect backsplicing. (7) Very rare AS-exonic circRNAs. (8) This is only our observation.

Based on the results of this study, we are convinced that the circularization *in-vitro* of RNA fragments during RNA preparation prior to sequencing is possible. The analyses conducted in this study revealed that a significant proportion of circRNAs identified in mRNAseq data were not detected in total-RNAseq data (63.4% for CD and 37.1% for CIRI2 analyses). These statistics would have been higher if sporadic circularization events had not been eliminated from our analyses. Moreover, we noticed that a similar high proportion of new circRNAs is often observed in datasets generated after RNase-R [5; 32]. For example, Gruhl et al (2021) [32] had to eliminate 75% of the circRNAs that were detected in RNase-R-treated samples but not in untreated samples. We believe that the partial digestion of linear RNAs by RNase-R contributes to an increase in the number of RNA fragments. This may be one of the reasons for the large number of other_circRNAs detected in 117T. The sub-exonic circRNAs originating from one exon of a multi-exonic gene, the circRNAs with their two genomic coordinates in two different exons of the same gene, and circRNAs from the mitochondrial genome are candidates to be artificial circRNAs with an *in-vivo* genesis. The common feature of sub-exonic circRNAs and circRNAs from the mitochondrial genome is that they originate from genes that are abundantly transcribed in the considered tissue [27]. We believe that the main feature of an artificial circRNAs obtained *in-vitro* is that this type of events is only weakly reproducible (to nucleotide precision) (Figure 11). This could happen e.g., via template switching during reverse transcription in the library preparation process [13; 53]. This *in-vitro* event does not lead to a circularization but only to the formation of a junction in the cDNA of junction that resembles to a circular junction. A genuine source of *in-vitro* circularization could be RNA fragments containing specific sequences that promote the formation of a double-stranded RNA with its two ends. The abundance of the initial linear transcript and treatments leading to RNA fragmentation probably increases the impact of such event. This is also the mechanism proposed by Liu et al. (2020) [26] for the genesis of interior circRNAs. Template switching may also be favored by the abundance of RNA fragments.

In contrast to many authors [6], only 20 antisense-exonic circRNAs were characterized in this study, while 23,737 sense-exonic circRNAs were characterized. It is important to note that we only considered the use of “reverse-complement exons” in theoretical backsplicing, which we termed AS-BS. When we examined the circular junction of these 20 circRNAs, we found that it is the mirror image of a previously characterized circular junction, which identifies a specific reliable exonic circRNA. Furthermore, such backsplicing involved “reverse-complement” exons is difficult to imagine. A strict-antisense transcript have an exon/intron structure that is symmetrical to another transcript of the parental gene. Since the “reverse-complement introns” do not have expected functional splicing sites at their boundaries (GT-AG becomes CT-AC), their origin cannot be explained by simple direct DNA transcription. They are likely generated by RNA polymerization using a spliced RNA as a template strand, i.e. by an RNA-dependent RNA polymerase (RdRP) [54; 55]. At first, we were very surprised to find that the lists of AS-exons (”reverse-complement exons”) involved in circRNAs and linear transcripts had nothing in common. In addition, this list of AS-exons involved in AS-BS was not the mirror image of the list of exons involved in the most common backsplicing events. These features suggest that these circRNAs are generated *in-vivo*. Nevertheless, these AS-exonic circRNAs cannot generated by a backsplicing event, and the circular nature of these RNAs has not been proven and is not obvious. The identification of AS-exonic circRNAs probably results from the detection of traces of (partial) copying of certain sense-exonic circRNAs, which would be generated *in-vivo* by an RdRP (Figure 11). These circRNAs are not detected by CIRI2 in 117T or by CD in 63m, which is consistent with this hypothesis.

On the Figure 11, we reported features of artificial circRNAs in comparison of exonic circRNAs. In addition to artificial circRNAs generated *in-silico* during the alignment process, *in-vitro* generation of artificial circRNAs should be considered. We noted that a new method has emerged recently to differentiate exonic circRNAs and other non-co-linear transcripts (fusion, trans-splicing) [56; 57]. Northern blotting is an interesting technique for revealing the circular configuration of RNA, but is rarely used [53; 58; 59]. A PCR amplification of the circular junction region as well as a test for resistance to RNase-R are often used to validate a circRNA [52; 53; 59]. We believe that only circular junctions generated *in-silico* after misalignment cannot be amplified by PCR whereas some *in-vitro* artificial circRNAs might pass these tests. We were not surprised to find in the literature that a significant fraction of non-exonic circRNAs detected by different tools could pass these validation tests [29]. Among the 1,516 circRNAs considered by Vromman et al (2023) [51], we found in the lists published by the authors at least 172 “other_circRNAs” that were validated by the three methods (qPCR, resistance to RNase-R and amplicon sequencing). Moreover, we identified 22 out of 39 sub-exonic circRNAs (circRNAs with both genomic coordinates in the same exon) from coding genes that were tested and were validated by the three methods. Using an approach focused on the *RPGRorf15* locus, Apelbaum et al. (2023) [60] confirmed the existence of several interior/sub-exonic circRNAs formed by back-fusion of linear parts (exonic and intronic) of the *RPGRorf15* pre-mRNAs. Further verification is required, but the current study suggests that the main feature of (sub-exonic) *in-vitro* artificial circRNAs may be the multiplicity of circRNAs from the same locus [27]. This is an easily detectable feature for highly expressed genes, but some circRNA tools tend to erase this feature. Another more surprising example is circRNAs from the mitochondrial genome, which, according to this study, are very likely to be artificial circRNAs [61].

This study showed that the number of bovine introns involved in intron circles was close to that involved in the production of lariat-derived circRNAs (ciRNAs), 126 and 147, respectively. These observations are in line with those recently made in humans [62]. This is not what has been observed previously in pigs, but that study involved only a very specific dataset [63]. Two genes are able to produce the two types of intronic circRNAs from distinct introns (ENSBTAG00000001888, *MED13L* and ENSBTAG00000032087, *ATXN2L*) but we found that they are not able to produce exonic circRNAs. The number of parental genes for intronic circRNAs (268) is significantly lower compared to exonic circRNAs (8 to 8.5K).

From the 117 tissue samples we analyzed, we found that the cerebellum was the tissue with the highest number of distinct exonic circRNAs in cattle. A similar result was observed in pigs [64]. We also found that the testis sampled from an adult animal could not be distinguished from the other tissues by the number of expressed exonic circRNAs. This result is consistent with comparisons made in pigs [27; 64]. This non-distinct clustering status of testis compared to other tissues with respect to the number of exonic circRNAs is somewhat surprising, as testis is highly transcriptionally active and is the tissue in which the highest number of protein-coding genes are expressed [65; 66]. Testicular exonic circRNAs seemed to be very tissue-specific, as demonstrated by the outlier status in the PCA analysis. These PCAs also showed that for the circular transcriptome there was some proximity between the adult adrenal and the adult testis and a large distance between these two tissues and the uterus, ovary and adrenal of non-pubertal animals. The circular transcriptome of the adrenal and testes is likely to be more affected by steroid synthesis than that of the ovary in bovine. However, this conclusion is probably due to the ovary sample used, which was taken from a cow three weeks after parturition, a period insufficient to observe normal ovarian function.

The overall tissue specificity of the circular transcriptome observed by hierarchical clustering analyses was very high for 8 tissues of 15 considered (kidney, cerebellum, muscle, heart, liver, lung, spleen, and adrenal gland). The 9th tissue for which we observed tissue-wise clustering of all samples was the rumen, but only when the data were normalized by the log-binary method. Clustering was biologically meaningful for two further tissues (fat and pituitary gland), if young animals are excluded from the analysis. No tissue specificity for the circular transcriptome was observed for four digestive tissues. Indeed, these five digestive tissues (duodenum, ileum, jejunum and colon) were the most resistant to clustering in the analyses of the 56 individual samples. These observations are not significantly different from those made in sheep considering only linear transcripts, which also showed that digestive tissues clustered poorly [67]. The results of the HCA constructed using the top-100 panel or the top-150 panel of expressed circRNA appear to be the most robust, yet construct with a panel containing 4.8 or 7.4% of the exonic circRNAs identified in these samples. It is quite surprising that we cannot improve the results of these HCA. However, these results again show that it is efficient to focus on highly expressed exonic circRNAs [27; 32].

## Conclusion

This study compared circRNAs present in 117 samples with total-RNAseq and mRNAseq data. Using this method, we confirmed the existence of several types of reliable circRNAs, including sense exonic circRNAs, ciRNA, intron circles, and sub-exonic circRNAs from snc genes. We highlighted the existence of 20 circRNAs, which are probably just traces of the copy of certain sense-exonic circRNAs. They would be generated *in-vivo* by an RNA-dependent RNA polymerase (RdRP). However, the study also identified a large number of circRNAs that are not generated *in-vivo.* The analysis of circRNA in mRNAseq datasets provided clear evidence that sub-exonic circRNAs from coding genes (introduced in [27]) are artifacts, while sub-exonic circRNAs from small non-coding genes (introduced in [28]) are not. Several hypotheses have been proposed to explain the presence of artif_circ RNAs in any RNAseq datasets. The most innovative are those related to *in-silico* and *in-vitro* factors. The possibility of *in-vitro* circularization of RNA fragments underlines the significance of the quality and integrity of the RNA source for the elaboration of datasets considered in circRNA studies. Our analysis leads us to recommend focusing on exonic circRNAs for tissue comparisons, such as those performed in this study of the bovine circular transcriptome for the BovReg project.

## Availability of data and materials

All data obtained concerning exonic and intronic circRNAs are available in the supplementary tables. The list of other_circRNAs is not available, as we were unable to distinguish between reliable and unreliable other_circRNAs. Datasets generated by the BovReg consortium and analyzed during the current study are listed in STab-1.

## Supplemental materials

### Materials and Methods, additional documents

MM&M_Adoc-1: exonic circRNA and backsplicing

M&M_Adoc-2: Circular RNAs: Vocabulary and annotations

M&M_Adoc-3: Example of region producing circRNAs

M&M_Adoc-4: genomic coordinates

M&M_Adoc-5: minimal_annotation of exonic circRNAs

M&M_Adoc-6: approach

### Results, Additional documents

Res_Adoc-1: Comparison of both annotations performed for the 23,926 exonic circRNAs

Res_Adoc-2: Overview across other_circRNAs detected in the different datasets and with alternative bioinformatic pipelines

Res_Adoc-3: Comparison of features of other_circRNAs

Res_Adoc-4: circRNA from *BTBD7*

Res_Adoc-5: circRNAs from *NEB*

Res_Adoc-6: Comparison 63T/63m (other_circRNAs)

Res_Adoc-7: Other_circRNAs identified in this study

Res_Adoc-8: large format of the Figure 7

Res_Adoc9: All results of HCAs

Res_Adoc-10: Overview of 61,083 circRNAs characterized in bovine tissues

**Supplementary. Tables** (not available in this version)

STab-1: List of samples considered

Stab-2: List of intronic circRNAs

STab-3: expression of 23,946 exonic circRNAs (sense and antisense) in 117 samples

STab-4: Presentation of 20 AS-exonic circRNA

STab-5: list of 189 circRNAs removed from the list of exonic circRNAs

STab-6: list of panels of exonic circRNAs (top-100 to top-250)

## Acknowledgments

Annie Robic acknowledges INRAE (more precisely the Animal Genetics and the M2I divisions) and the FBN (more precisely the Institute of Genome Biology), which supported her research stay in FBN in 2022-2023. We thank Dr Sylvain Foissac and Dr Laurence Liaubet (GenPhySE) for indirectly enriching this study through their insightful discussions.

## M&M_Adoc-1

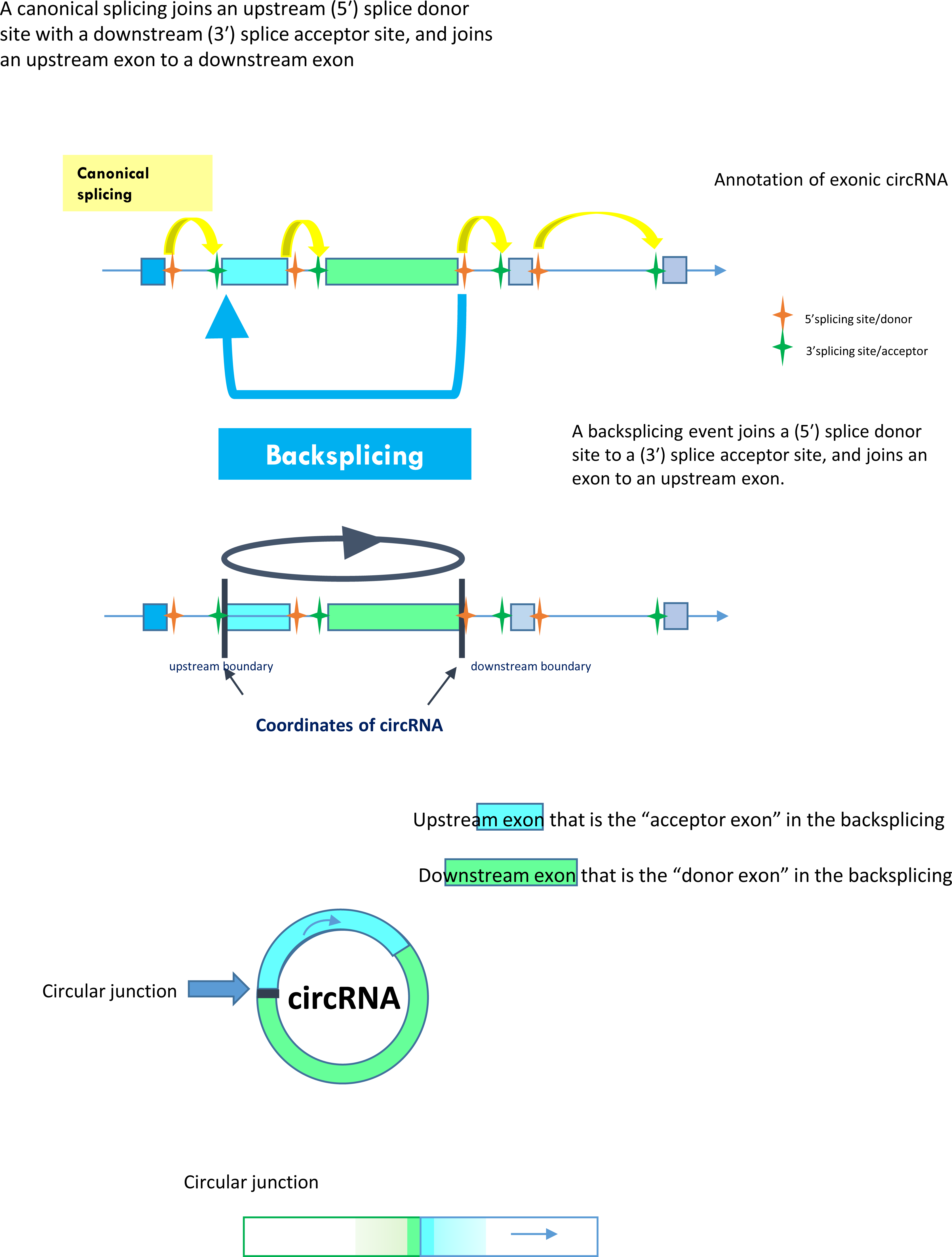

## M&M_Adoc-2

### Circular RNAs : Vocabulary and annotations

➢ ***A few vocabulary questions***

- Total-RNAseq: sequence data from total RNA libraries after depletion of ribosomal RNA
- mRNA-Seq: sequencing data from mRNA libraries, i.e. after polyA selection
- circRNA: to design all types of circular RNA supported by reads spanning a circular junction. These reads are aligned to the reference genome sequence as two segments mapping in reverse order.
- Those annotated as “ciRNA” correspond to circRNAs localized entirely in intronic sequences and with the circRNA 5′ junction site corresponding to the intron donor site
- Intronic circRNA: intron circle + lariat derived circRNA
- Exonic circRNAs: only circRNAs resulting from backsplicing between two known exons. See also Suppl. Figure S1
- As we considered all circRNAs, not just those derived from backsplicing, it appeared to us that the term BSJ (backsplicing junction) is not appropriate to name the circular junction.
➢ ***Presentations of CD*** The source code of the CD is available from https://github.com/GenEpi-GenPhySE/circRNA.git. CD identifies reads containing a circular junction within those reads that STAR calls “chimeric reads” (CR) from the tabular file (chimeric.out.junction) provided by STAR: Subsequently, these reads will be called “circular chimeric reads” (CCRs). The main CD-output file (detection.bed) consists of a list of all circular RNAs and their associated number of CCRs, each circular RNA is being defined by the coordinates of the circular junction (chromosome:start-end|strand). For the detection step, the user can select a threshold x to retain only circRNAs characterized by at least x CCRs and a minimal genomic size for circRNAs (distance between the two feet of the circRNA). In this study, we have chosen not to consider non-redundant or sporadic circularization events. Several studies have shown the value of excluding such events [Gruhl et al. 2021 ; Xu et al. 2021]. The detection of circRNAs was performed individually for each dataset by CD with a threshold of 5 CCRs. The annotation is made to the precision of the base. CD does not tolerate any differences from the gtf file and does not perform any grouping. If there are three circRNAs, after annotation, it will keep three. There is one parameter that can be modified to enable grouping, but we strongly advise against using it. This circRNA tool was specifically designed for intronic circRNA characterization [Robic et al. 2022 ; Robic et al. 2021]. The identification of the reads supporting the circular junction is performed from the single end alignments of the PE reads, and the compatibility of the second read with this characterization is not verified. Given the difficulty of sequencing the 2’-3’ circular junction, this is an undeniable advantage for the characterization of intronic circRNAs [Robic et al. 2020a ; Robic et al. 2020b].
➢ ***Annotation of exonic circRNAs*** Both junctions correspond to exonic boundaries from a single gene located on the same strand as circRNA. Consequently, the circRNAs must satisfy the three following rules

- The 3’ junction of a circRNA must precisely correspond to an exon donor site (3’ end of an exon, ie 5’ donor site of the next intron) from a gene located on the same strand as circRNA
- The 5’ junction must precisely correspond to an upstream exon acceptor site (5’ end of an exon, ie 3’ acceptor site of the previous intron) from a gene located on the same strand as circRNA
- The exon donor and the exon acceptor are associated to a common gene
➢ ***Annotation of intronic circRNAs: lariat-derived intronic circRNA (or ciRNA) and intron circle***

- both junctions are located within a single intron
- the 5’ junction must precisely correspond to the 5’ intron donor site
- the 3’ junction must be compatible with a circularization event limited by the branch point (12-60 base pair away from the 3’ intron acceptor site) for lariat derived circRNAs
- the 3’ junction must be compatible with a circularization of the entire intron (-5/5 base pair away from the 3’ intron acceptor site) for intron circle
➢ ***Annotation of sub-exonic circRNAs from snc genes***

- Both junctions are located within a single exon
- This class contains no circRNA classified as Exonic
- Only the ones that are associated to a gene not reported as lnc, coding gene or pseudo-gene

**M&M_Adoc-3.**
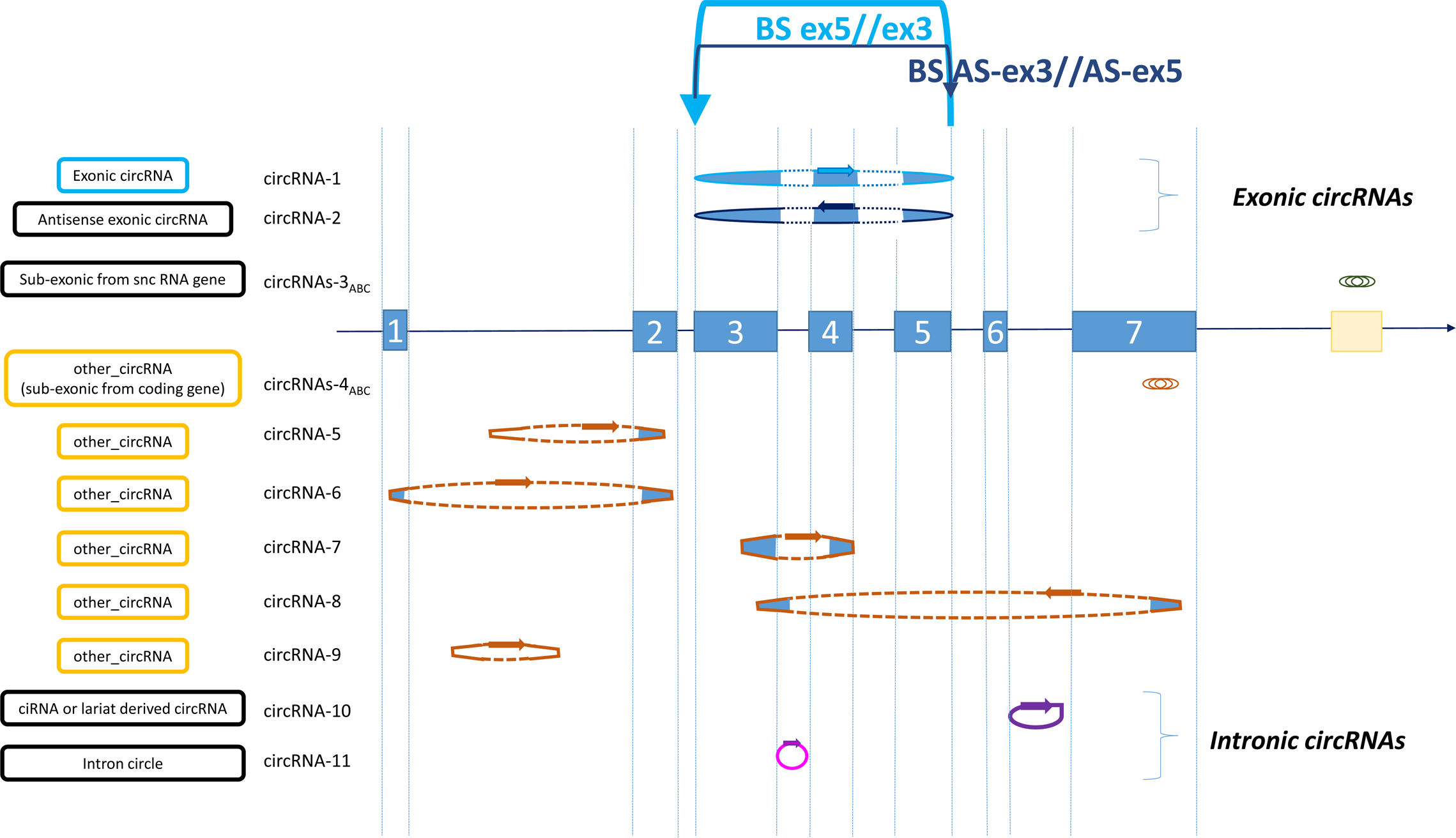
The circRNA-1 is an exonic circRNA and the circRNA-2 is an antisense exonic circRNA from the blue gene. The set of circRNAs-3 groups sub-exonic circRNAs from a snc gene and they can be in sense or antisense. Only circRNA-10 and -11 are intronic circRNAs (purple and pink). The first is a lariat-derived circRNA (ciRNA) and the second is an intron circle and they contain only intronic sequences. We chose to classify those that are not identified by CD as being exonic or intronic circRNAs or sub-exonic circRNAs originating from the snc gene as other_circRNAs. It is sort of a provisional class and our goal is to sort it out. circRNA-5, -6, -7, -8 and -9 and the set of circRNAs-4ABC are other_circRNAs (in brown). The classification exonic/other_cirRNA/miscellaneous appears as a colored frame (blue/orange/black) on the left.

## M&M_Adoc-4

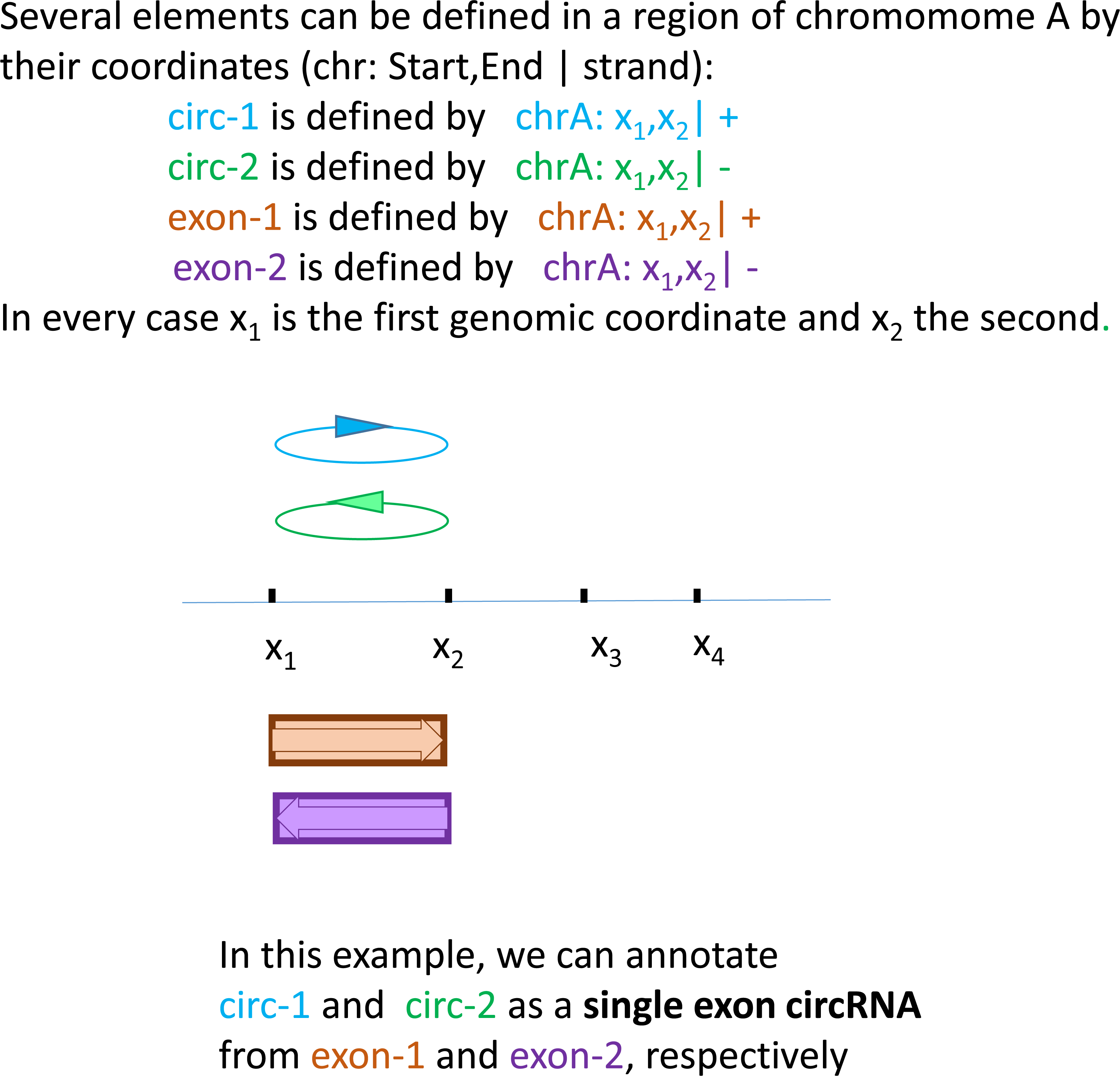

## M&M_Adoc-5A

**The constitution of the two sub-lists** to perform the minimal annotation

**To perform what we call a minimal_annotation of exonic circRNAs, we created two sub-lists (Left_exons and Right_exons) from the list of all BovReg exons.**

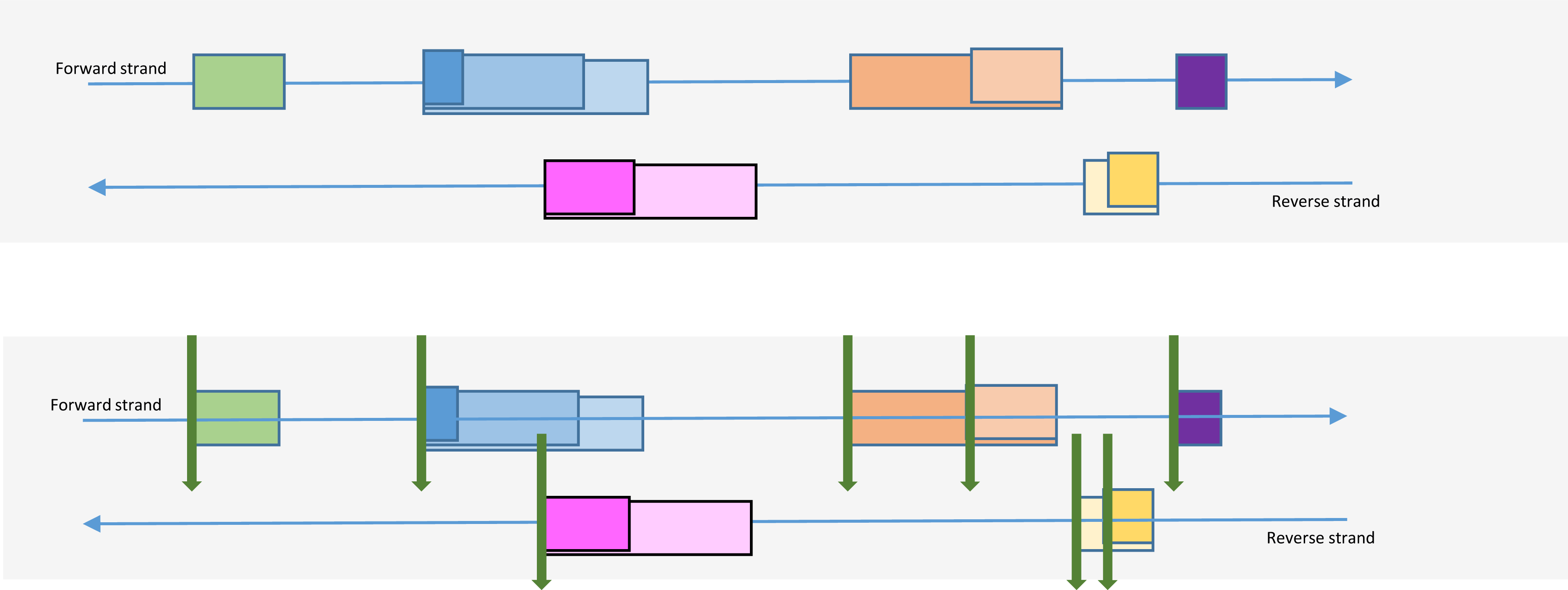

To constitute the list of Left_exons, we selected exons according to their unique first genomic coordinate keeping only the exon with the smallest size in case of multiple exons with the same first coordinate. This first coordinate (indicate by a green arrow) corresponds to the 5’ coordinate when the exon is defined on forward strand and to the 3’ coordinate for exons defined on reverse strand.

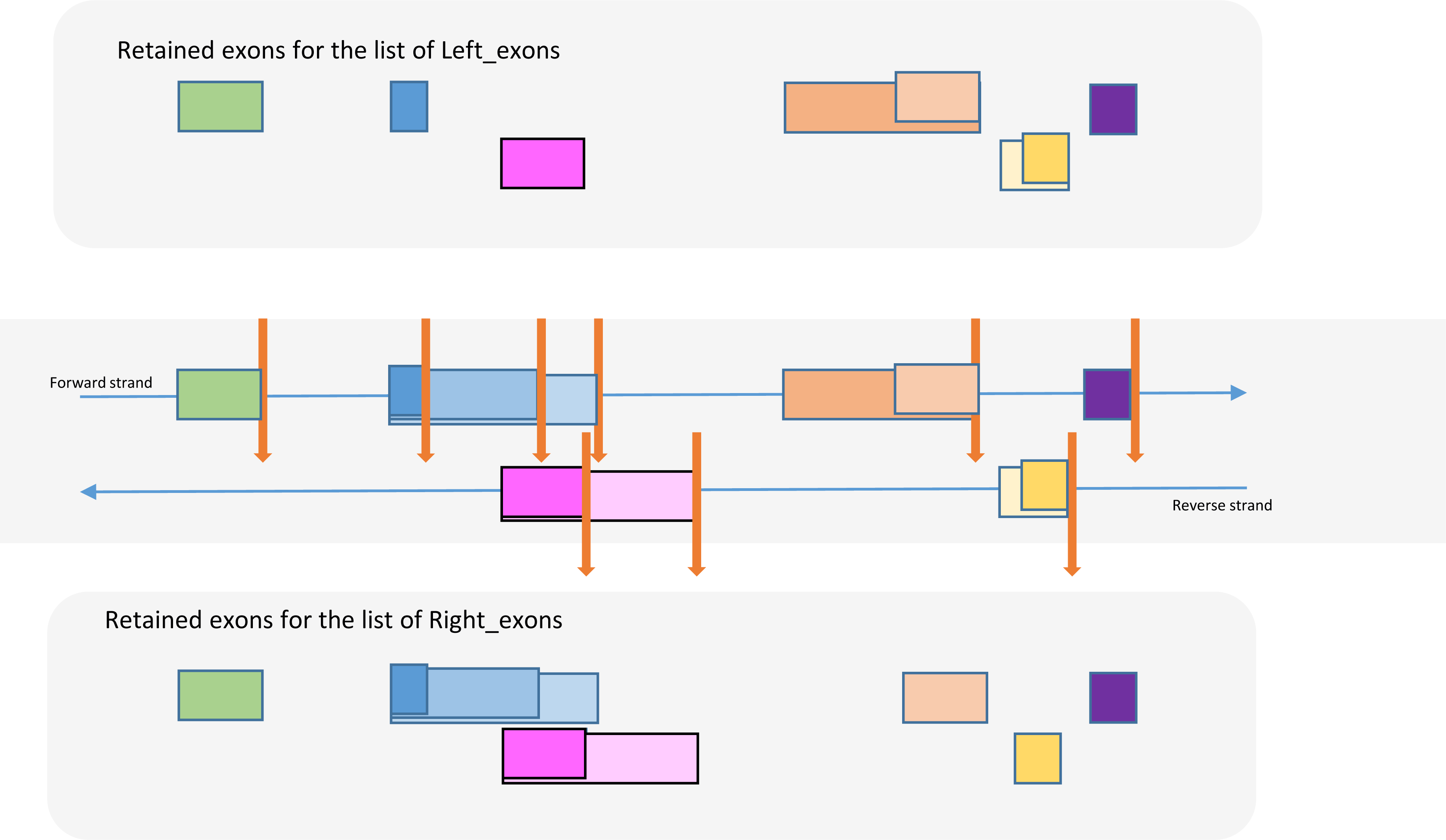

## M&M_Adoc-5B

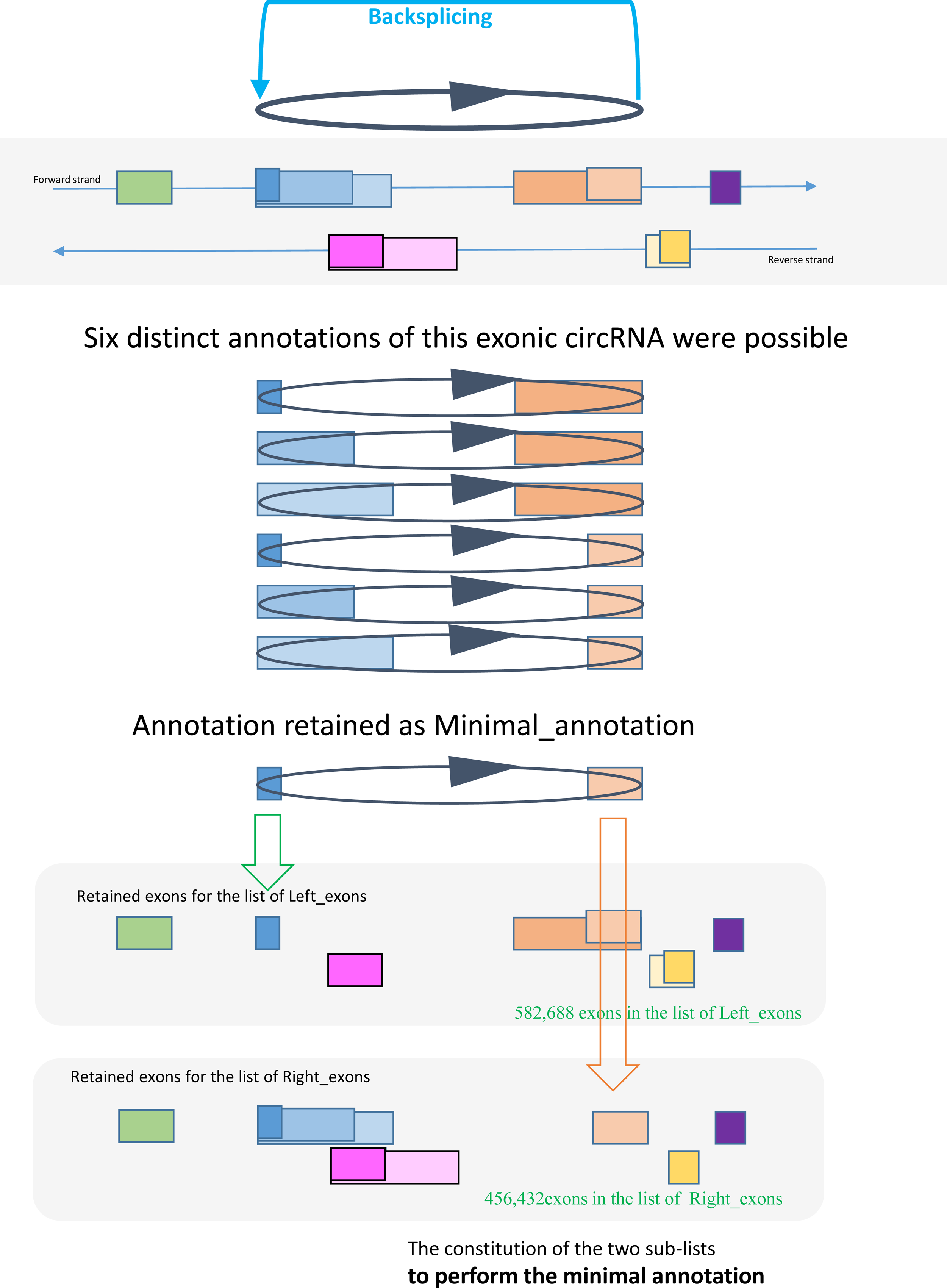

## M&M_Adoc-6

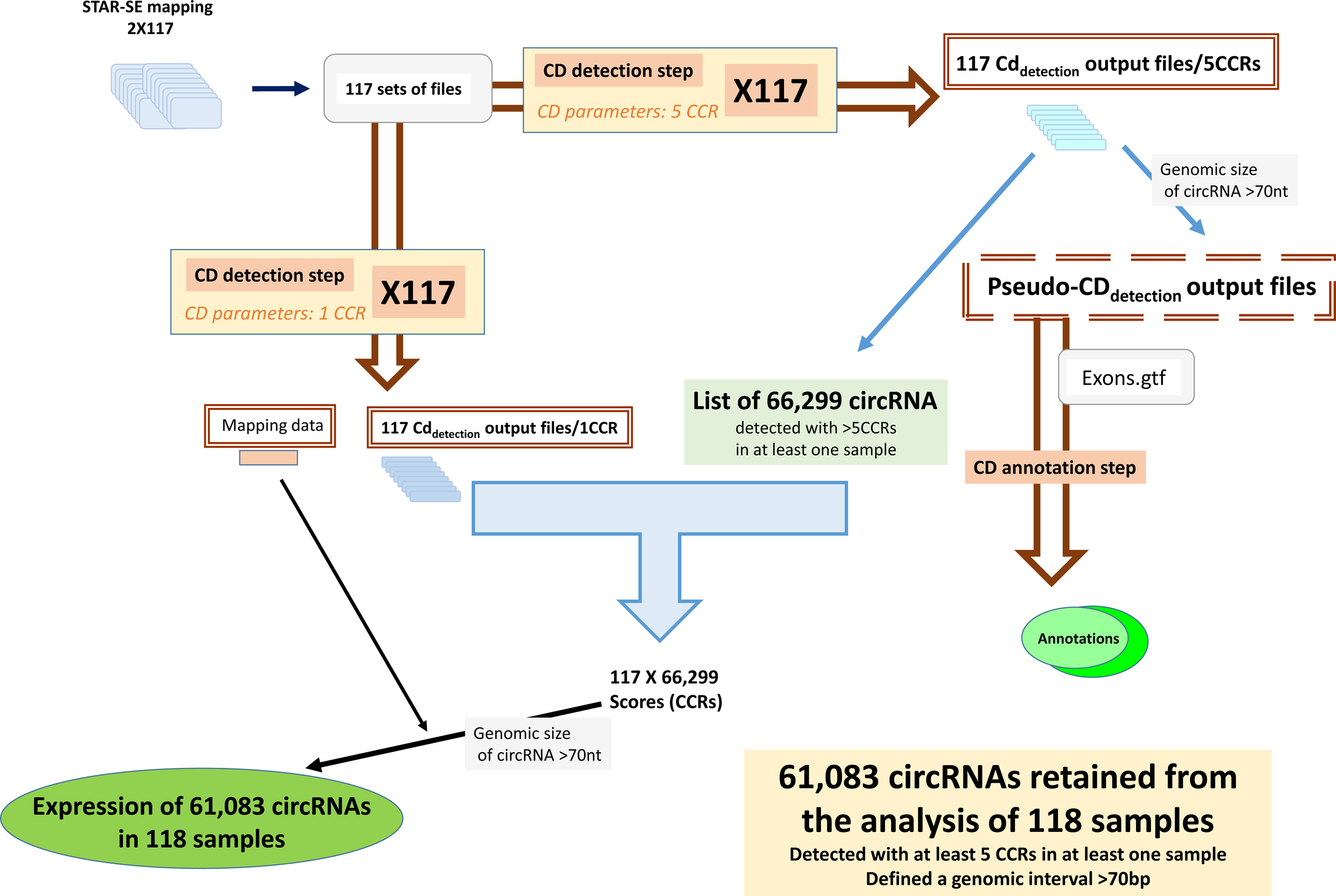

**Res_Adoc-1.** Comparison of both annotations performed for the 23,926 exonic circRNAs.

| Classical annotation<br>(CD + manual curation) | Exon | Minimal annotation |
| --- | --- | --- |
| 20,000 | Ens-Ens | 18,429 |
| 1,894 | MSTRG-MSTRG | 1,682 |
| 2,032 | Mixed<br>(Ens-MSTRG or MSTRG-Ens) | 3,815 |
| 7,993 | Number of parental genes | 8,461 |

**Res_Adoc-1.** Overview across other_circRNAs detected in the different datasets and with alternative bioinformatic pipelines.

Res\_Adoc-2 Overview across other\_circRNAs detected in the different datasets and with alternative bioinformatic pipelines
| Sample | circRNAs considered | Detected | with size < 160nt |  |
| --- | --- | --- | --- | --- |
| 117T | All CD-other_circRNAs | 36,215 | 8,285 | 22.88% |
|  | *Defensin* region | 3,159 | 403 | 12.80% |
|  | Mitochondrial genome | 487 | 235 | 48.30% |
| 63T | All CD-other_circRNAs | 17,956 | 3,966 | 22.10% |
| 63m | All CD-other_circRNAs | 4,341 | 2,129 | (49,0%) |
| CIRI-117T | All CIRI-other_circRNAs | 10,081 | 111 | 1.10% |

Three comparisons of the number of other\_circRNAs with small size were tested >> chisq\_test with p-value < 2.2 10<sup>-16</sup>

**Res_Adoc-3.** Comparison of features of other_circRNAs.

|  | CD-other_circRNAs | CIRI-other_circRNAs |
| --- | --- | --- |
| Two feet out of any exon | 29.90% | 18.00% |
| Only one foot in exonic sequences | 7.70% | 32.70% |
| Two feet in exonic sequences | 62.40% | 49.30% |
| The interval defined by the two feet contains a full exon | 28.80% | 74.00% |
| The size of the genomic interval is >500 kb | 1% | 0%* |
| The size of the genomic interval is <160 bp | 22.88% | 1.10% |

\* CIRI2 does not retain circRNAs defining a genomic interval >200 kb
We defined the two ‘feet’ of a circRNA by the two genomic coordinates of its genomic boundaries.

## Res_Adoc-4

### Exonic circRNAs from *BTBD7*

21:57913322-57915074|-

The recently published study of an exonic circRNA from the bovine *BTBD7* gene led us to focus on this gene with 11 exons and a length of 85.6 kb (ENSBTAG00000046185) [1]. We have characterized only one exonic circRNA produced by this gene. This is the one characterized by Ma et al. (2023) and it was detected in 117/117 samples.

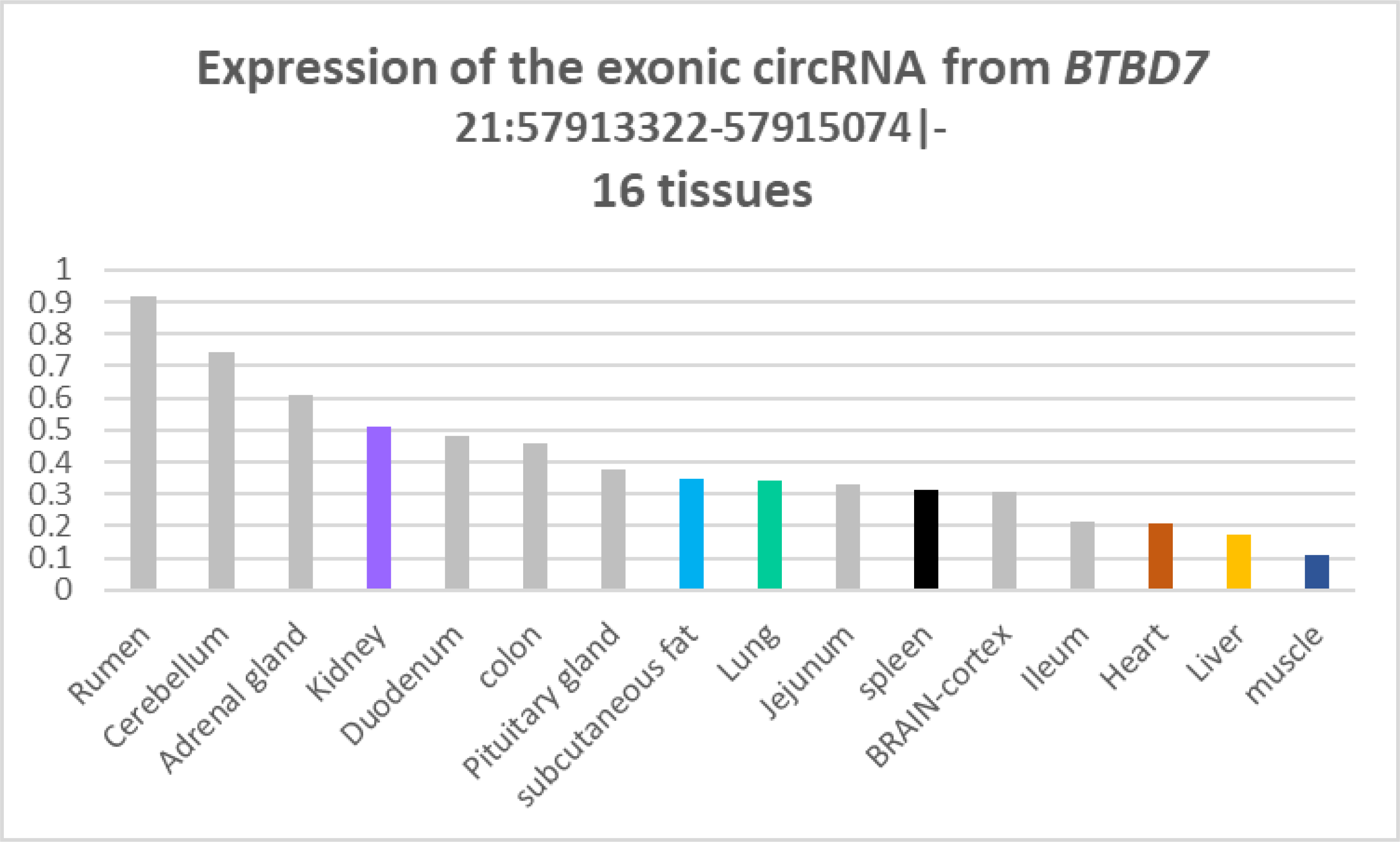

The 16 tissues were ranked in descending order of average expression.

(for the BRAIN-cortex we considered the both datasets obtained from the juvenile animal (castrated male)

The colors used are identical to those used by Ma et al. (2023) in their Figure 2E.

We observed that the highest expression of this exonic RNA is not found in subcutaneous fat,

But in rumen. Nevertheless, our observations on tissue expression of this exonic circRNA are almost all compatible with the results published by Ma et al. (2023). If we examine the expression of this exonic circRNAs in the 5 or 65 samples available for some tissues, we note that individual variability is significant.

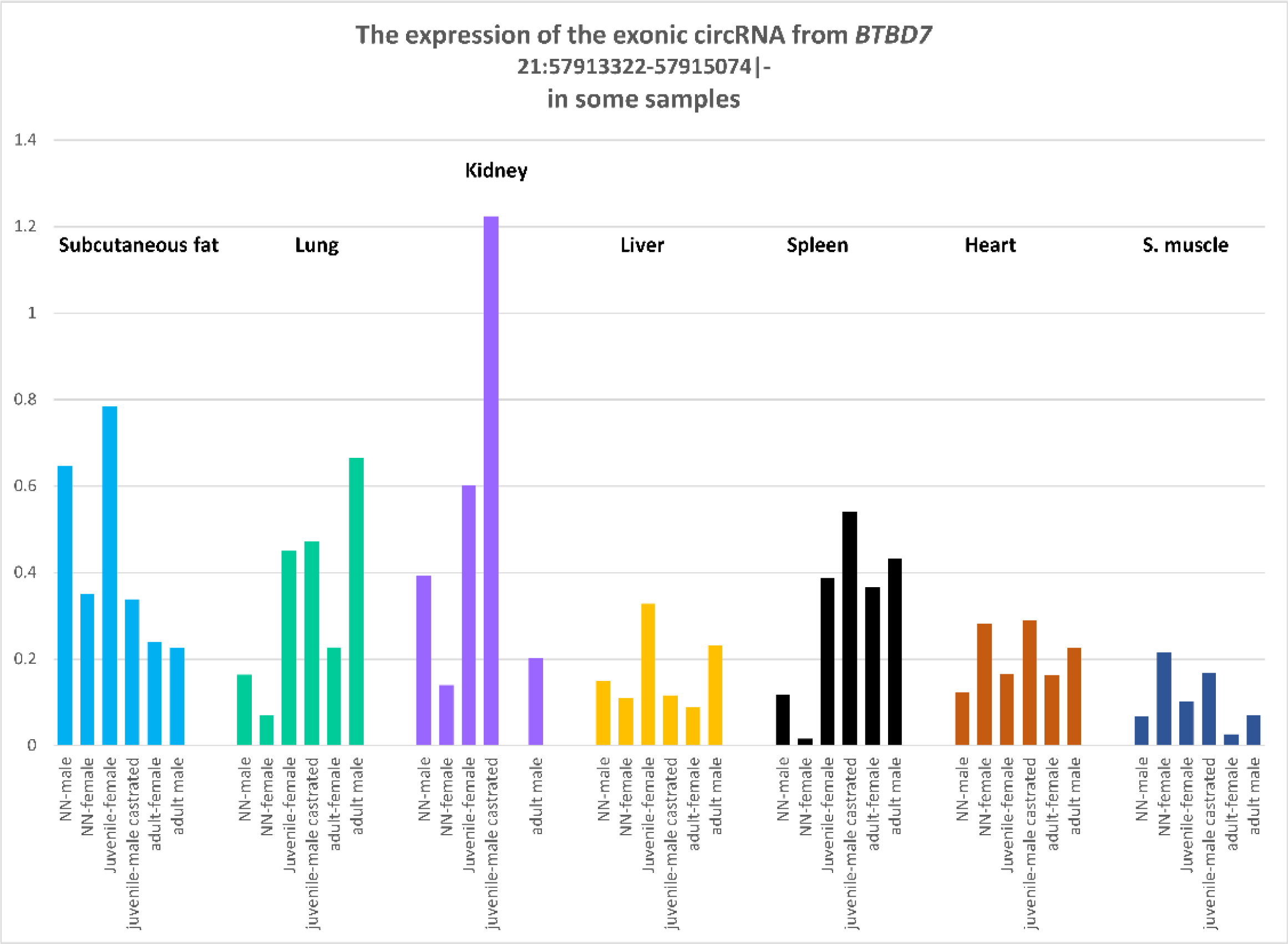

## Res_Adoc-5

### Exonic circRNAs from *NEB*

A recent study published on a specific bovine exonic circRNA [2] has brought attention to a particular gene. The bovine *Nebulin* gene (ENSBTAG00000006907, NEB) gene is a very large gene (219 kb) with more than 170 exons in which we observe several repetitions of four exons: five repetitions of the exonic sequence A (exons 204-105-108-312 nt, 243 aa) followed by twelve repetitions of the exonic sequence B (exons 207-105-108-312 nt, 244 aa). Huang et al. (2024) [2] characterized an exonic circRNA with four exons (204, 105, 108 and 312 nt) encoding for a rolling-translated peptide. With these features, it was not surprising to observe that the rolling-translated sequence of the circNEB polypeptide is highly homologous to the partial sequence of the NEB protein [2]. Out of the 23,926 sense exonic circRNAs detected in this study, the circNEB characterized by Huang et al. was not found. More surprisingly, none of the exonic circRNAs characterized in this study were identified as surrogates. Given the limited sample size of only six muscle samples out of the 117 studied, further examination of published results is recommended. It is worth noting that among the bovine exonic circRNAs characterized in a study including 12 muscle samples from adults animals (and 15 liver and 6 testis) using CD [3], we found the same absence. On the contrary, when we considered the list of circRNAs characterized by CircExplorer2+CIRI2 [3], we found three exonic circRNAs compatible with a rolling-translated peptide (with two, three or four repetitions of the exonic sequence B).

---

**Res_Adoc-6.**
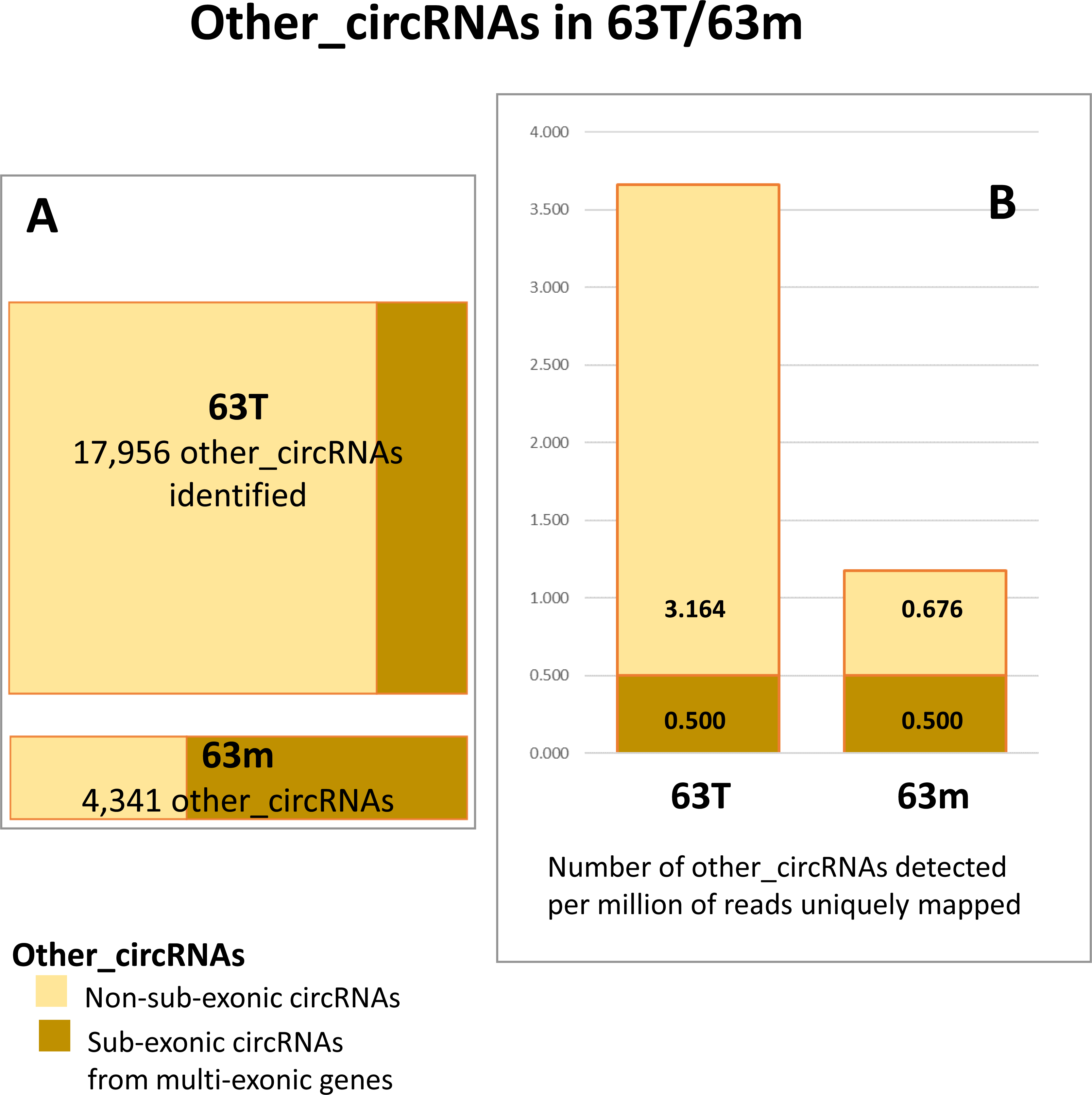

**Res_Adoc-7.** Other_circRNAs identified in this study.

Other\_circRNAs identified in this study
| Number of other_circRNA identified |  | 63m | 63T | 118T |
| --- | --- | --- | --- | --- |
| **Genome** |  | **4,341** | **17,956** | **36,245** |
|  | already known | 1,529 | 17,956 | 36,245 |
|  | novel | 2,812 |  |  |
| **Albumin region** |  | **27** | **59** | **59** |
|  | already known | 22 | 59 | 59 |
|  | novel | 5 |  |  |
| **Defensin region** |  | **7** | **1,671** | **3,160** |
| BTA27: 6,21-6,23Mb | already known | 4 | 1,671 | 3,160 |
|  | novel | 3 |  |  |
| **MT chromosome** |  | **556** | **319** | **487** |
|  | already known | 98 | 319 | 487 |
|  | novel | 458 |  |  |

**Res_Adoc-8A.**
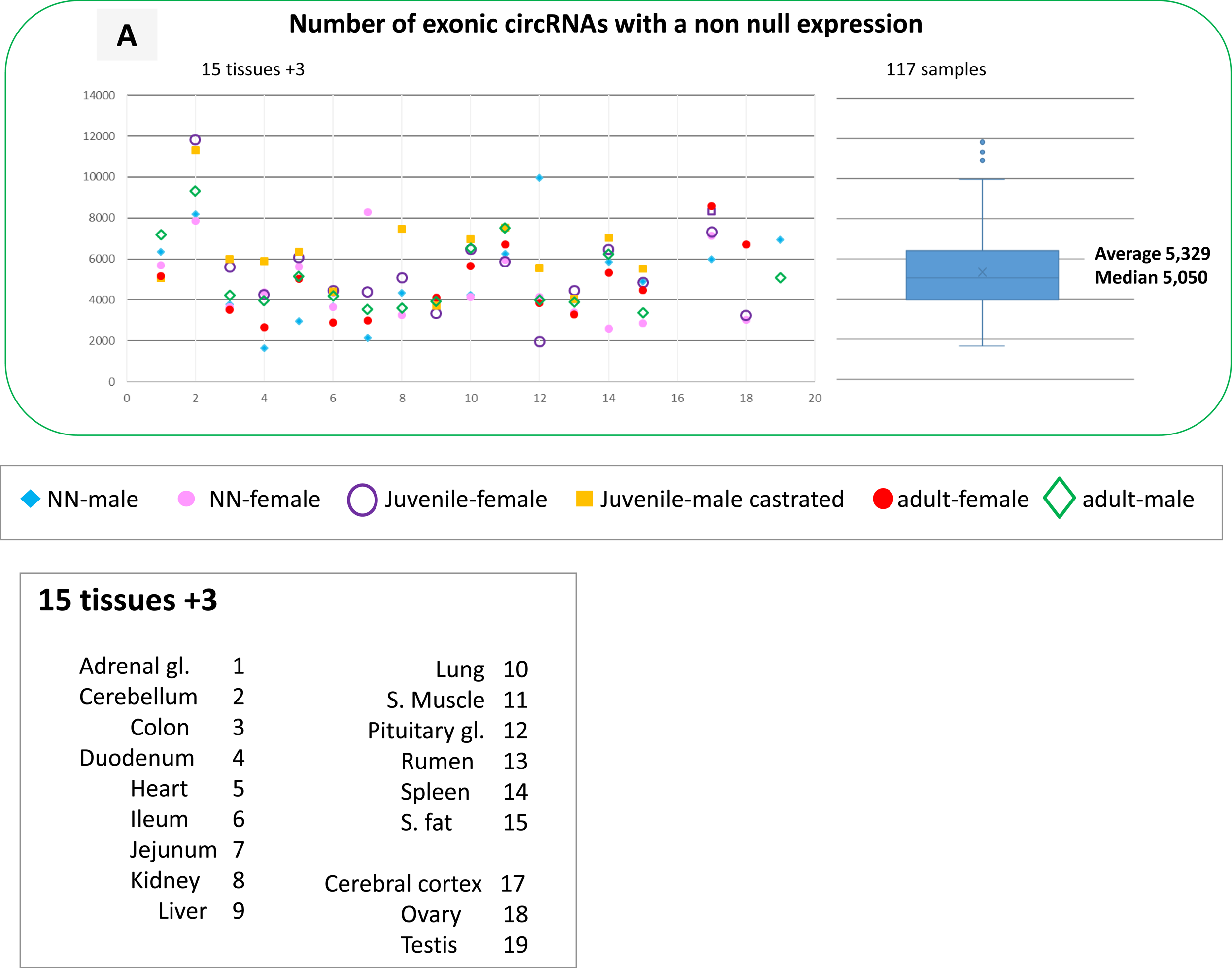

**Res_Adoc-8B.**
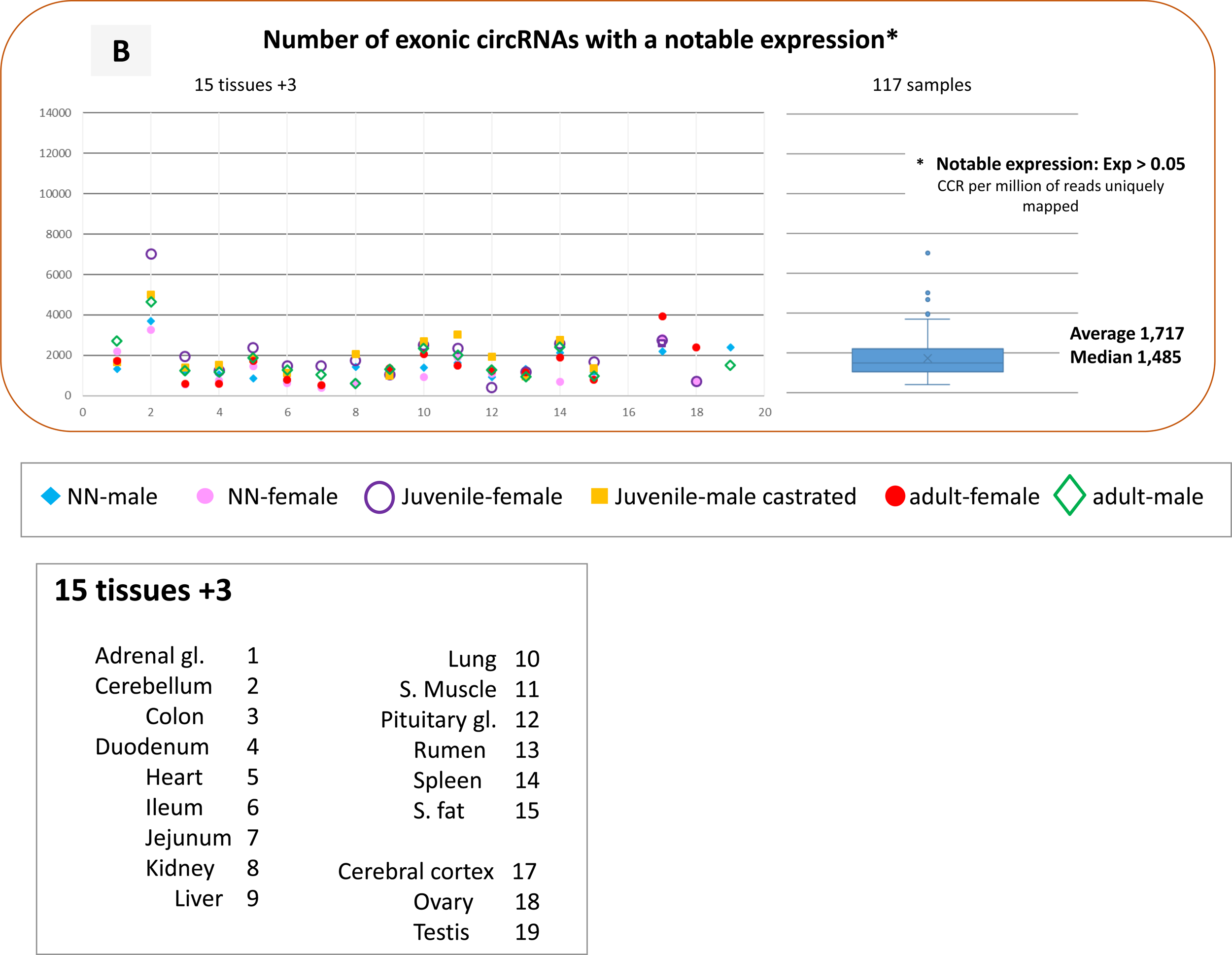

**Res_Adoc-8C.**
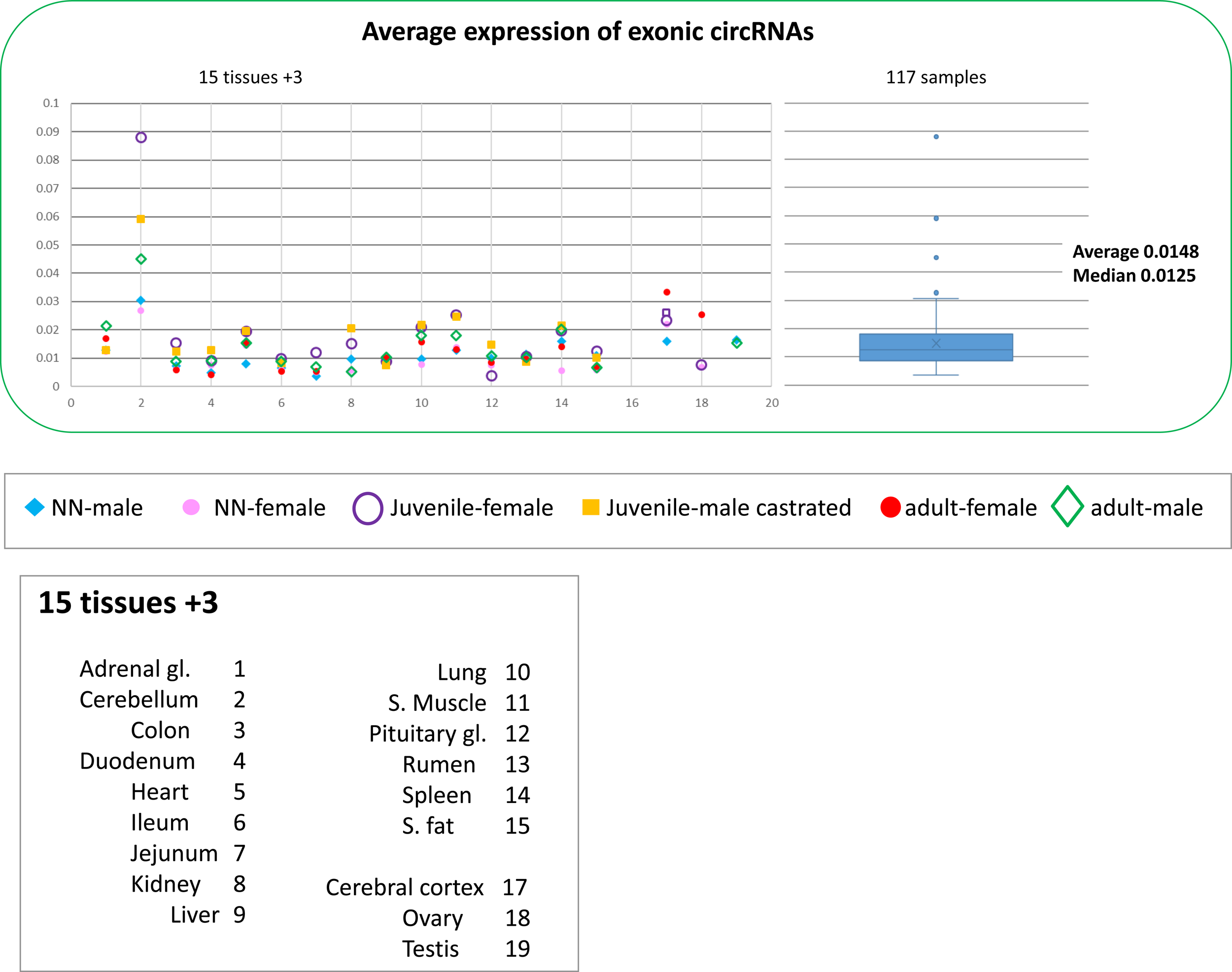

**Res_Adoc-9.**
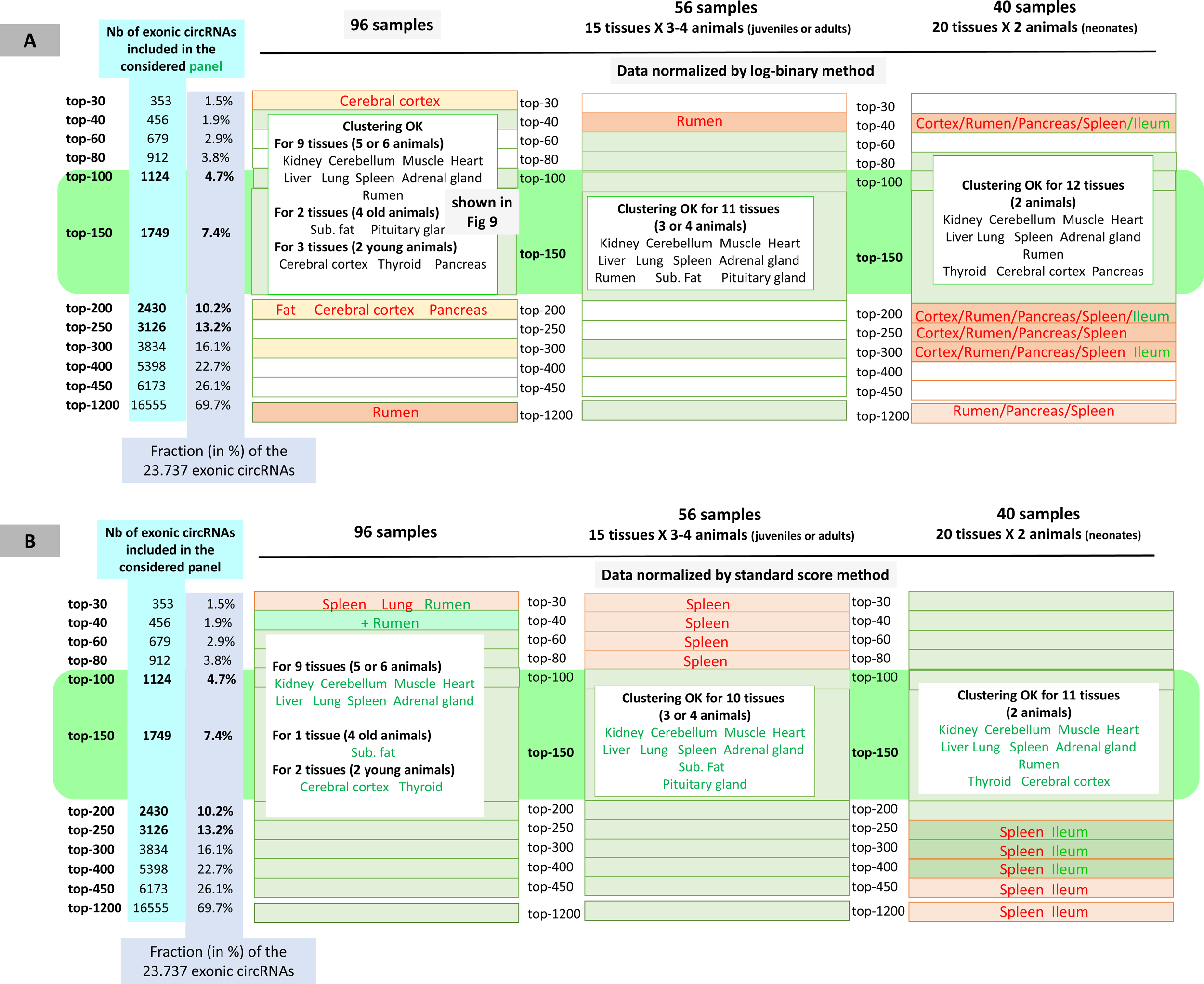
All HCAs were built using the “ward” agglomeration method and Pearson correlations as distance. Results of all HCAs built using different panels of exonic circRNAs with data normalized by the log-binary method (B) or by the standard score (C) were summarized. Only the observations that differed from the respective reference results (HCA obtained with top-150) were reported, From left to right, 96 samples from 20 tissues were considered, only the 56 samples from oldest animals (J and A) were considered, and only the 40 samples from neonatal animals (N) were considered.

**Res_Adoc-10.**
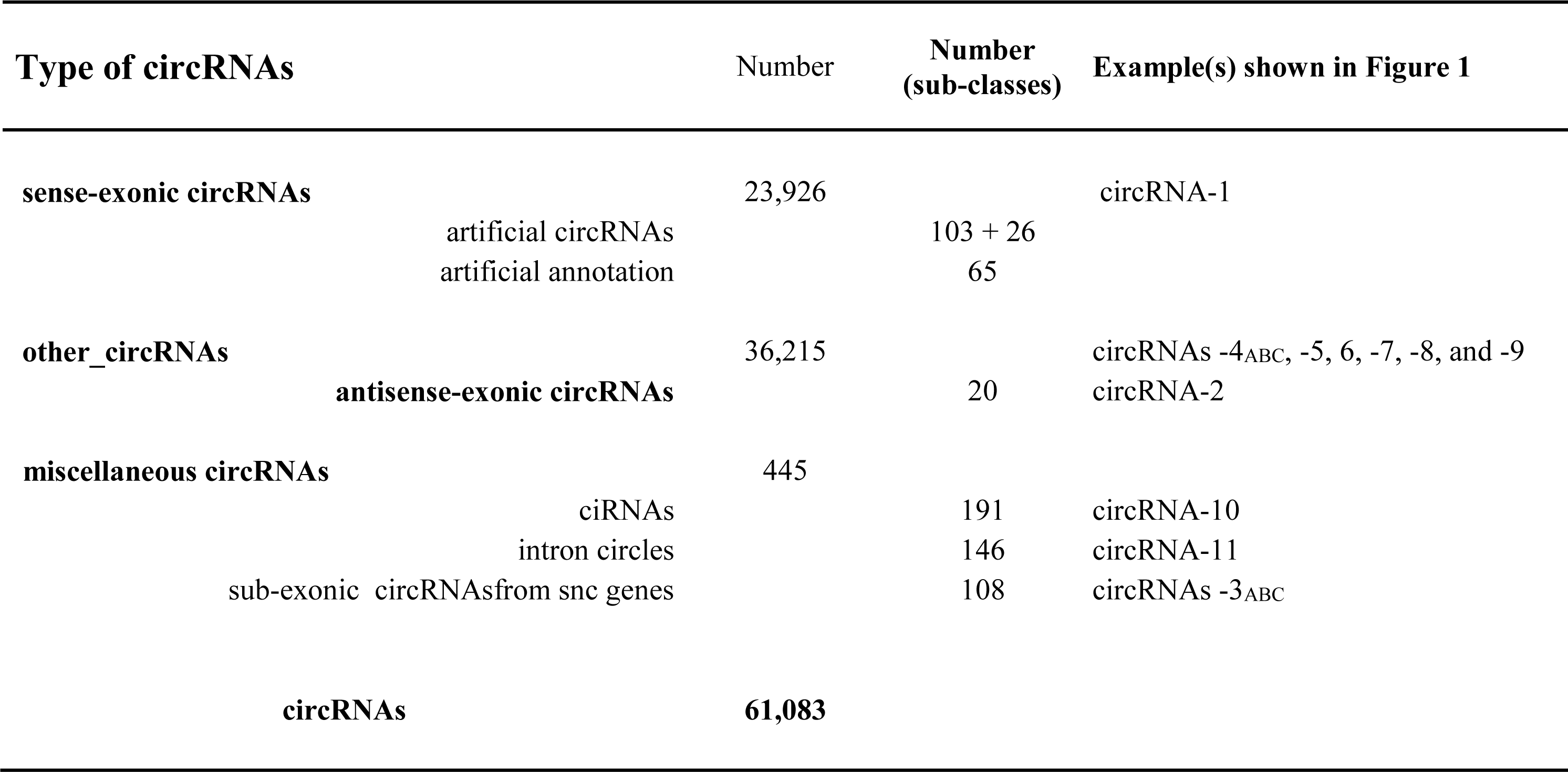
Overview of 61,083 circRNAs characterized in bovine tissues.

**Why the AS-exonic circRNAs are found in other_circRNAs category?**

These AS-exonic circRNAs involve BS with AS-exons. Some exons “transcribed in A” have been identified in linear transcript studies but none of those involved in BS. For this reason, we found ASexonic circRNAs in the list of other_circRNAs.

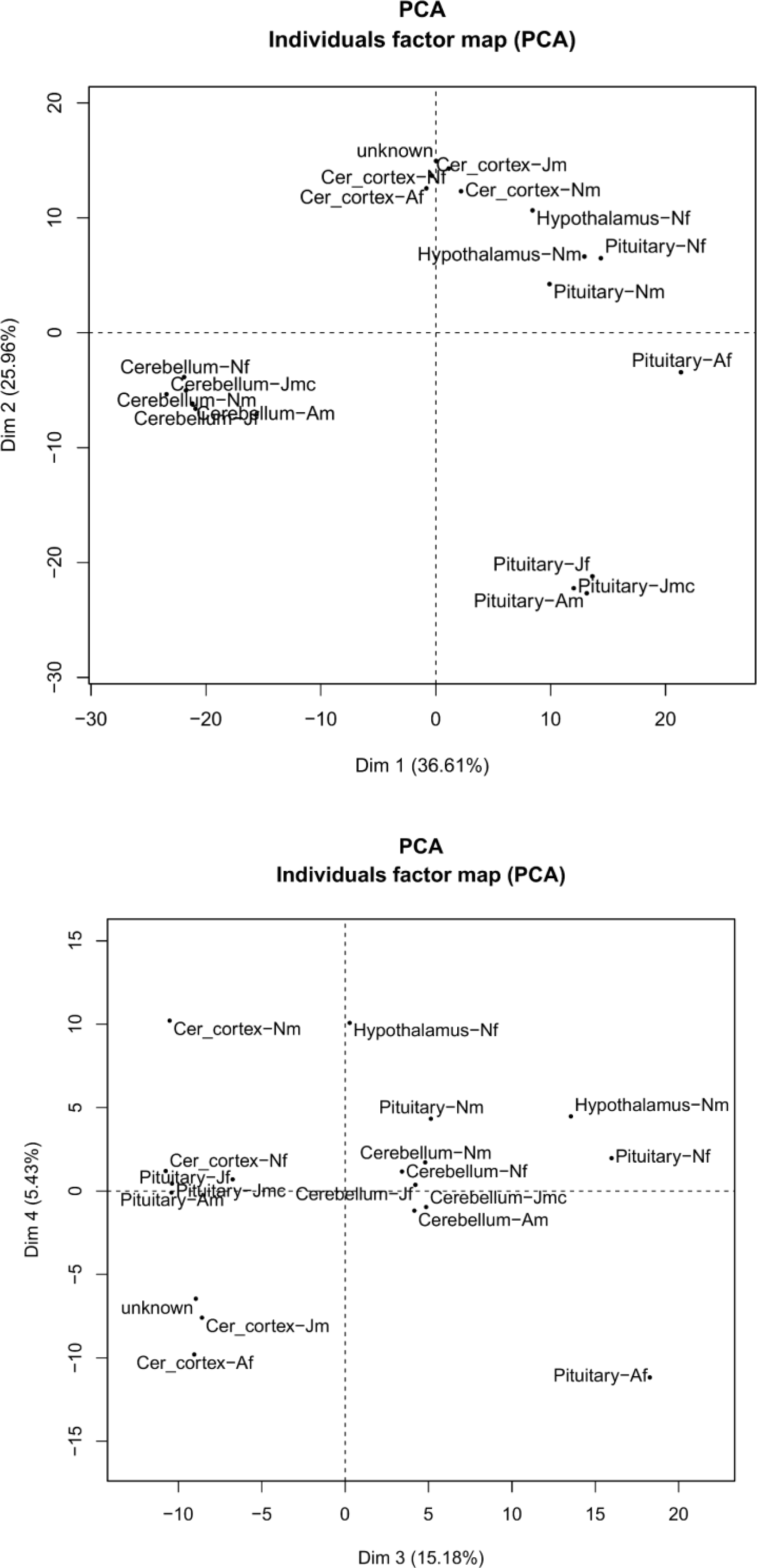

## REFERENCES

[1] B.D. Rosen, D.M. Bickhart, R.D. Schnabel, S. Koren, C.G. Elsik, E. Tseng, T.N. Rowan, W.Y. Low, A. Zimin, C. Couldrey, R. Hall, W. Li, A. Rhie, J. Ghurye, S.D. McKay, F. Thibaud-Nissen, J. Hoffman, B.M. Murdoch, W.M. Snelling, T.G. McDaneld, J.A. Hammond, J.C. Schwartz, W. Nandolo, D.E. Hagen, C. Dreischer, S.J. Schultheiss, S.G. Schroeder, A.M. Phillippy, J.B. Cole, C.P. Van Tassell, G. Liu, T.P.L. Smith, and J.F. Medrano, De novo assembly of the cattle reference genome with single-molecule sequencing. GigaScience 9 (2020).

[2] D.E. Goszczynski, M.M. Halstead, A.D. Islas-Trejo, H. Zhou, and P.J. Ross, Transcription initiation mapping in 31 bovine tissues reveals complex promoter activity, pervasive transcription, and tissue-specific promoter usage. Genome Res 31 (2021) 732–744.

[3] E.M. Ross, H. Sanjana, L.T. Nguyen, Y. Cheng, S.S. Moore, and B.J. Hayes, Extensive Variation in Gene Expression is Revealed in 13 Fertility-Related Genes Using RNA-Seq, ISO-Seq, and CAGE-Seq From Brahman Cattle. Frontiers in genetics 13 (2022) 784663.

[4] M. Salavati, R. Clark, D. Becker, C. Kuhn, G. Plastow, S. Dupont, G.C.M. Moreira, C. Charlier, and E.L. Clark, Improving the annotation of the cattle genome by annotating transcription start sites in a diverse set of tissues and populations using Cap Analysis Gene Expression sequencing. G3 13 (2023).

[5] W.R. Jeck, J.A. Sorrentino, K. Wang, M.K. Slevin, C.E. Burd, J. Liu, W.F. Marzluff, and N.E. Sharpless, Circular RNAs are abundant, conserved, and associated with ALU repeats. RNA 19 (2013) 141–57.

[6] S. Memczak, M. Jens, A. Elefsinioti, F. Torti, J. Krueger, A. Rybak, L. Maier, S.D. Mackowiak, L.H. Gregersen, M. Munschauer, A. Loewer, U. Ziebold, M. Landthaler, C. Kocks, F. le Noble, and N. Rajewsky, Circular RNAs are a large class of animal RNAs with regulatory potency. Nature 495 (2013) 333–8.

[7] J. Salzman, C. Gawad, P.L. Wang, N. Lacayo, and P.O. Brown, Circular RNAs are the predominant transcript isoform from hundreds of human genes in diverse cell types. PLoS One 7 (2012) e30733.

[8] Y. Zhang, X.O. Zhang, T. Chen, J.F. Xiang, Q.F. Yin, Y.H. Xing, S. Zhu, L. Yang, and L.L. Chen, Circular intronic long noncoding RNAs. Molecular cell 51 (2013) 792–806.

[9] L.S. Kristensen, M.S. Andersen, L.V.W. Stagsted, K.K. Ebbesen, T.B. Hansen, and J. Kjems, The biogenesis, biology and characterization of circular RNAs. Nat Rev Genet 20 (2019) 675–691.

[10] C.X. Liu, and L.L. Chen, Circular RNAs: Characterization, cellular roles, and applications. Cell 185 (2022) 2390.

[11] J.E. Wilusz, Circular RNAs: Unexpected outputs of many protein-coding genes. RNA biology 14 (2017) 1007–1017.

[12] L. Yang, J.E. Wilusz, and L.L. Chen, Biogenesis and Regulatory Roles of Circular RNAs. Annual review of cell and developmental biology 38 (2022) 263–289.

[13] T. Horiuchi, and T. Aigaki, Alternative trans-splicing: a novel mode of pre-mRNA processing. Biology of the cell 98 (2006) 135–40.

[14] T.J. Chuang, Y.J. Chen, C.Y. Chen, T.L. Mai, Y.D. Wang, C.S. Yeh, M.Y. Yang, Y.T. Hsiao, T.H. Chang, T.C. Kuo, H.H. Cho, C.N. Shen, H.C. Kuo, M.Y. Lu, Y.H. Chen, S.C. Hsieh, and T.W. Chiang, Integrative transcriptome sequencing reveals extensive alternative trans-splicing and cis-backsplicing in human cells. Nucleic Acids Res (2018).

[15] J. Dubois, and G. Sczakiel, The human TRAM1 locus expresses circular RNAs. Scientific reports 11 (2021) 22114.

[16] K. Rahimi, M.T. Veno, D.M. Dupont, and J. Kjems, Nanopore sequencing of brain-derived full-length circRNAs reveals circRNA-specific exon usage, intron retention and microexons. Nature communications 12 (2021) 4825.

[17] A. Robic, T. Faraut, S. Djebali, R. Weikard, K. Feve, S. Maman, and C. Kuehn, Analysis of pig transcriptomes suggests a global regulation mechanism enabling temporary bursts of circular RNAs. RNA biology 16 (2019) 1190–1204.

[18] G.J. Talhouarne, and J.G. Gall, Lariat intronic RNAs in the cytoplasm of Xenopus tropicalis oocytes. RNA 20 (2014) 1476–87.

[19] A.J. Taggart, C.L. Lin, B. Shrestha, C. Heintzelman, S. Kim, and W.G. Fairbrother, Large-scale analysis of branchpoint usage across species and cell lines. Genome Res 27 (2017) 639–649.

[20] G.J.S. Talhouarne, and J.G. Gall, Lariat intronic RNAs in the cytoplasm of vertebrate cells. Proc Natl Acad Sci U S A 115 (2018) E7970–E7977.

[21] X.K. Ma, S.N. Zhai, and L. Yang, Approaches and challenges in genome-wide circular RNA identification and quantification. Trends Genet (2023).

[22] A.F. Nielsen, A. Bindereif, I. Bozzoni, M. Hanan, T.B. Hansen, M. Irimia, S. Kadener, L.S. Kristensen, I. Legnini, M. Morlando, M.T. Jarlstad Olesen, R.J. Pasterkamp, S. Preibisch, N. Rajewsky, C. Suenkel, and J. Kjems, Best practice standards for circular RNA research. Nature methods 19 (2022) 1208–1220.

[23] X.K. Ma, W. Xue, L.L. Chen, and L. Yang, CIRCexplorer pipelines for circRNA annotation and quantification from non-polyadenylated RNA-seq datasets. Methods 196 (2021) 3–10.

[24] Y. Gao, J. Zhang, and F. Zhao, Circular RNA identification based on multiple seed matching. Brief Bioinform 19 (2018) 803–810.

[25] X. Liu, J. Frost, A. Bowcock, and W. Zhang, Canonical and Interior Circular RNAs Function as Competing Endogenous RNAs in Psoriatic Skin. Int J Mol Sci 22 (2021).

[26] X. Liu, Z. Hu, J. Zhou, C. Tian, G. Tian, M. He, L. Gao, L. Chen, T. Li, H. Peng, and W. Zhang, Interior circular RNA. RNA biology 17 (2020) 87–97.

[27] A. Robic, C. Cerutti, C. Kühn, and T. Faraut, Comparative analysis of the circular transcriptome in muscle, liver and testis in three livestock species. Frontiers in genetics 12 (2021) 665153.

[28] A. Robic, J. Demars, and C. Kühn, In-Depth Analysis Reveals Production of Circular RNAs from Non-Coding Sequences. Cells 9 (2020) 1806.

[29] C.Y. Yu, H.J. Liu, L.Y. Hung, H.C. Kuo, and T.J. Chuang, Is an observed non-co-linear RNA product spliced in trans, in cis or just in vitro? Nucleic Acids Res 42 (2014) 9410–23.

[30] X. Lv, W. Chen, W. Sun, Z. Hussain, L. Chen, S. Wang, and J. Wang, Expression profile analysis to identify circular RNA expression signatures in hair follicle of Hu sheep lambskin. Genomics 112 (2020) 4454–4462.

[31] T. Lu, L. Cui, Y. Zhou, C. Zhu, D. Fan, H. Gong, Q. Zhao, C. Zhou, Y. Zhao, D. Lu, J. Luo, Y. Wang, Q. Tian, Q. Feng, T. Huang, and B. Han, Transcriptome-wide investigation of circular RNAs in rice. RNA 21 (2015) 2076–87.

[32] F. Gruhl, P. Janich, H. Kaessmann, and D. Gatfield, Circular RNA repertoires are associated with evolutionarily young transposable elements. Elife 10 (2021).

[33] G.C.M. Moreira, S. Dupont, D. Becker, M. Salavati, R. Clark, E.L. Clark, G. Plastow, C. Kühn, and C. Charlier, Multi-dimensional functional annotation of the bovine genome for the BovReg project. Proceedings of 12th World Congress on Genetics Applied to Livestock Production (WCGALP). (2022) 2261–2264.

[34] A. Dobin, C.A. Davis, F. Schlesinger, J. Drenkow, C. Zaleski, S. Jha, P. Batut, M. Chaisson, and T.R. Gingeras, STAR: ultrafast universal RNA-seq aligner. Bioinformatics 29 (2013) 15–21.

[35] J. Cheng, F. Metge, and C. Dieterich, Specific identification and quantification of circular RNAs from sequencing data. Bioinformatics 32 (2016) 1094–6.

[36] A. Robic, C. Cerutti, J. Demars, and C. Kuhn, From the comparative study of a circRNA originating from an mammalian ATXN2L intron to understanding the genesis of intron lariat-derived circRNAs. Biochimica et biophysica acta. Gene regulatory mechanisms 1865 (2022) 194815.

[37] H. Li, Toward better understanding of artifacts in variant calling from high-coverage samples. Bioinformatics 30 (2014) 2843–51.

[38] SIGENAE, http://www.sigenae.org/.

[39] R core team, R: A language and environment for statistical computing. Foundation for Statistical Computing, Vienna, Austria (2020).

[40] HCA-Galaxy-tutorial, http://genoweb.toulouse.inra.fr/~formation/CATIBIOS4BIOL_stats/Learning_clustering_current.pdf.

[41] C. Xu, and J. Zhang, Mammalian circular RNAs result largely from splicing errors. Cell reports 36 (2021) 109439.

[42] L.L. Chen, A. Bindereif, I. Bozzoni, H.Y. Chang, A.G. Matera, M. Gorospe, T.B. Hansen, J. Kjems, X.K. Ma, J.W. Pek, N. Rajewsky, J. Salzman, J.E. Wilusz, L. Yang, and F. Zhao, A guide to naming eukaryotic circular RNAs. Nature cell biology 25 (2023) 1–5.

[43] C. Xue, J. Wei, M. Li, S. Chen, L. Zheng, Y. Zhan, Y. Duan, H. Deng, W. Xiong, G. Li, H. Li, and M. Zhou, The Emerging Roles and Clinical Potential of circSMARCA5 in Cancer. Cells 11 (2022).

[44] Z. Ma, Y. Chen, J. Qiu, R. Guo, K. Cai, Y. Zheng, Y. Zhang, X. Li, L. Zan, and A. Li, CircBTBD7 inhibits adipogenesis via the miR-183/SMAD4 axis. International journal of biological macromolecules 253 (2023) 126740.

[45] D. Liu, B.K. Dredge, A.G. Bert, K.A. Pillman, J. Toubia, W. Guo, B.J.A. Dyakov, M.M. Migault, V.M. Conn, S.J. Conn, P.A. Gregory, A.C. Gingras, D. Patel, B. Wu, and G.J. Goodall, ESRP1 controls biogenesis and function of a large abundant multiexon circRNA. Nucleic Acids Res (2023).

[46] K. Huang, Z. Li, D. Zhong, Y. Yang, X. Yan, T. Feng, X. Wang, L. Zhang, X. Shen, M. Chen, X. Luo, K. Cui, J. Huang, S.U. Rehman, Y. Jiang, D. Shi, A. Pauciullo, X. Tang, Q. Liu, and H. Li, A Circular RNA Generated from Nebulin (NEB) Gene Splicing Promotes Skeletal Muscle Myogenesis in Cattle as Detected by a Multi-Omics Approach. Advanced science 11 (2024) e2300702.

[47] C. Ragan, G.J. Goodall, N.E. Shirokikh, and T. Preiss, Insights into the biogenesis and potential functions of exonic circular RNA. Scientific reports 9 (2019) 2048.

[48] G. Santos-Rodriguez, I. Voineagu, and R.J. Weatheritt, Evolutionary dynamics of circular RNAs in primates. Elife 10 (2021).

[49] S. Starke, I. Jost, O. Rossbach, T. Schneider, S. Schreiner, L.H. Hung, and A. Bindereif, Exon circularization requires canonical splice signals. Cell reports 10 (2015) 103–11.

[50] L. Szabo, R. Morey, N.J. Palpant, P.L. Wang, N. Afari, C. Jiang, M.M. Parast, C.E. Murry, L.C. Laurent, and J. Salzman, Statistically based splicing detection reveals neural enrichment and tissue-specific induction of circular RNA during human fetal development. Genome Biol 16 (2015) 126.

[51] M. Vromman, J. Anckaert, S. Bortoluzzi, A. Buratin, C.Y. Chen, Q. Chu, T.J. Chuang, R. Dehghannasiri, C. Dieterich, X. Dong, P. Flicek, E. Gaffo, W. Gu, C. He, S. Hoffmann, O. Izuogu, M.S. Jackson, T. Jakobi, E.C. Lai, J. Nuytens, J. Salzman, M. Santibanez-Koref, P. Stadler, O. Thas, E. Vanden Eynde, K. Verniers, G. Wen, J. Westholm, L. Yang, C.Y. Ye, N. Yigit, G.H. Yuan, J. Zhang, F. Zhao, J. Vandesompele, and P.J. Volders, Large-scale benchmarking of circRNA detection tools reveals large differences in sensitivity but not in precision. Nature methods 20 (2023) 1159–1169.

[52] T.J. Chuang, T.W. Chiang, and C.Y. Chen, Assessing the impacts of various factors on circular RNA reliability. Life science alliance 6 (2023) e202201793.

[53] S. Dodbele, N. Mutlu, and J.E. Wilusz, Best practices to ensure robust investigation of circular RNAs: pitfalls and tips. EMBO Rep 22 (2021) e52072.

[54] S.D. Wagner, P. Yakovchuk, B. Gilman, S.L. Ponicsan, L.F. Drullinger, J.F. Kugel, and J.A. Goodrich, RNA polymerase II acts as an RNA-dependent RNA polymerase to extend and destabilize a non-coding RNA. The EMBO journal 32 (2013) 781–90.

[55] Y. Maida, and K. Masutomi, RNA-dependent RNA polymerases in RNA silencing. Biological chemistry 392 (2011) 299–304.

[56] T.J. Chuang, C.S. Wu, C.Y. Chen, L.Y. Hung, T.W. Chiang, and M.Y. Yang, NCLscan: accurate identification of non-co-linear transcripts (fusion, trans-splicing and circular RNA) with a good balance between sensitivity and precision. Nucleic Acids Res 44 (2016) e29.

[57] Y.C. Chen, C.Y. Chen, T.W. Chiang, M.H. Chan, M. Hsiao, H.M. Ke, I.J. Tsai, and T.J. Chuang, Detecting intragenic trans-splicing events from non-co-linearly spliced junctions by hybrid sequencing. Nucleic Acids Res 51 (2023) 7777–7797.

[58] T. Schneider, S. Schreiner, C. Preusser, A. Bindereif, and O. Rossbach, Northern Blot Analysis of Circular RNAs. Methods in molecular biology 1724 (2018) 119–133.

[59] Z. Mi, C. Zhongqiang, J. Caiyun, L. Yanan, W. Jianhua, and L. Liang, Circular RNA detection methods: A minireview. Talanta 238 (2022) 123066.

[60] T. Appelbaum, G.D. Aguirre, and W.A. Beltran, Identification of circular RNAs hosted by the RPGR ORF15 genomic locus. RNA biology 20 (2023) 31–47.

[61] Z. Wu, H. Sun, C. Wang, W. Liu, M. Liu, Y. Zhu, W. Xu, H. Jin, and J. Li, Mitochondrial Genome-Derived circRNA mc-COX2 Functions as an Oncogene in Chronic Lymphocytic Leukemia. Molecular therapy. Nucleic acids 20 (2020) 801–811.

[62] A.M. Rasmussen, T.L.H. Okholm, M. Knudsen, S. Vang, L. Dyrskjot, T.B. Hansen, and J.S. Pedersen, Circular stable intronic RNAs possess distinct biological features and are deregulated in bladder cancer. NAR cancer 5 (2023) zcad041.

[63] A. Robic, and C. Kühn, Beyond Back Splicing, a Still Poorly Explored World: Non-Canonical Circular RNAs. Genes 11 (2020) 1111.

[64] L. Jin, Q. Tang, S. Hu, Z. Chen, X. Zhou, B. Zeng, Y. Wang, M. He, Y. Li, L. Gui, L. Shen, K. Long, J. Ma, X. Wang, Z. Chen, Y. Jiang, G. Tang, L. Zhu, F. Liu, B. Zhang, Z. Huang, G. Li, D. Li, V.N. Gladyshev, J. Yin, Y. Gu, X. Li, and M. Li, A pig BodyMap transcriptome reveals diverse tissue physiologies and evolutionary dynamics of transcription. Nature communications 12 (2021) 3715.

[65] M. Soumillon, A. Necsulea, M. Weier, D. Brawand, X. Zhang, H. Gu, P. Barthes, M. Kokkinaki, S. Nef, A. Gnirke, M. Dym, B. de Massy, T.S. Mikkelsen, and H. Kaessmann, Cellular source and mechanisms of high transcriptome complexity in the mammalian testis. Cell reports 3 (2013) 2179–90.

[66] W. Yang, F. Zhao, M. Chen, Y. Li, X. Lan, R. Yang, and C. Pan, Identification and characterization of male reproduction-related genes in pig (Sus scrofa) using transcriptome analysis. BMC Genomics 21 (2020) 381.

[67] E.L. Clark, S.J. Bush, M.E.B. McCulloch, I.L. Farquhar, R. Young, L. Lefevre, C. Pridans, H.G. Tsang, C. Wu, C. Afrasiabi, M. Watson, C.B. Whitelaw, T.C. Freeman, K.M. Summers, A.L. Archibald, and D.A. Hume, A high resolution atlas of gene expression in the domestic sheep (Ovis aries). PLoS Genet 13 (2017) e1006997.

[68] M. Ares, H. Igel, S. Katzman, J. P. Donohue, Intron-lariat spliceosomes convert lariats to true circles: implications for intron transposition. bioRxiv 2024.03.26.586863; doi: 10.1101/2024.03.26.586863

## References

[1] Z. Ma, Y. Chen, J. Qiu, R. Guo, K. Cai, Y. Zheng, Y. Zhang, X. Li, L. Zan, and A. Li, CircBTBD7 inhibits adipogenesis via the miR-183/SMAD4 axis. International journal of biological macromolecules 253 (2023) 126740.

[2] K. Huang, Z. Li, D. Zhong, Y. Yang, X. Yan, T. Feng, X. Wang, L. Zhang, X. Shen, M. Chen, X. Luo, K. Cui, J. Huang, S.U. Rehman, Y. Jiang, D. Shi, A. Pauciullo, X. Tang, Q. Liu, and H. Li, A Circular RNA Generated from Nebulin (NEB) Gene Splicing Promotes Skeletal Muscle Myogenesis in Cattle as Detected by a Multi-Omics Approach. Advanced science 11 (2024) e2300702.

[3] A. Robic, C. Cerutti, C. Kühn, and T. Faraut, Comparative analysis of the circular transcriptome in muscle, liver and testis in three livestock species. Frontiers in genetics 12 (2021) 665153.

